# Combinatorial effects of *Zingiber officinale* and *Citrus limon* juices: Hypolipidemic and antioxidant insights from *in vivo*, *in vitro*, and *in silico* investigations

**DOI:** 10.1101/2025.02.08.637272

**Authors:** Oussama Bekkouch, Ayoub Bekkouch, Hamza Elbouny, Ilham Touiss, Soufiane El Assri, Mohammed Choukri, Mohammed Bourhia, Kiryang Kim, Sojin Kang, Min Choi, Jinwon Choi, Hyo Jeong Kim, Chi-Hoon Ahn, Moon Nyeo Park, Bonglee Kim, Souliman Amrani

## Abstract

The combined use of *Zingiber officinale* (ginger) and *Citrus limon* (lemon) has long been valued in traditional medicine for their significant therapeutic properties, particularly in supporting cardiovascular health. However, their synergistic effects on hyperlipidemia and oxidative stress, both critical risk factors for cardiovascular diseases, remain underexplored.

This study employed a multidisciplinary approach including *in vivo* (mice model with triton WR-1339-induced hyperlipidemia), *in vitro* (antioxidant assays such as DPPH and ABTS), and *in silico* (molecular docking targeting HMG-CoA reductase) to evaluate the hypolipidemic and antioxidant potential of ginger and lemon juices. Phytochemical analysis identified high concentrations of bioactive compounds, with ginger juice containing 6-gingerol (15.54% peak area) and lemon juice featuring hesperidin (13.85% peak area) and rutin (5.57% peak area). These compounds were instrumental in the observed pharmacological effects.

The combined formulation significantly improved lipid profiles, reducing total cholesterol by 54.3%, triglycerides by 49.8%, and LDL by 58.1% while increasing HDL by 47.6% compared to hyperlipidemic controls (p < 0.05). Antioxidant assays revealed high free radical scavenging activity, with IC_50_ values of 30.92 ± 2.00 µg/mL for ABTS and 44.94 ± 1.02 µg/mL for DPPH in the combined formulation. Molecular docking confirmed strong binding affinities, with hesperidin and 6-gingerol showing binding energies of -10.4 kcal/mol and -9.6 kcal/mol, respectively, against HMG-CoA reductase.

In conclusion, combining *Zingiber officinale* and *Citrus limon* juices demonstrates robust hypolipidemic and antioxidant effects, significantly outperforming the individual juices. These results support their potential as natural therapeutic agents for managing hyperlipidemia and oxidative stress. Further clinical studies are recommended to validate these promising findings and explore their application in nutraceutical development.

## 1. Introduction

Lipids play a crucial role in regulating numerous cellular physiological functions, and changes in membrane lipid metabolism are linked to significant diseases such as cancer, type II diabetes, cardiovascular disease, and immune disorders (Cockcroft, 2021); however, an elevated presence of plasmatic fats heightens the susceptibility to coronary heart disease. Dyslipidemia is a spectrum of abnormalities in lipoproteins and lipids, a primary risk factor for atherosclerosis (Farkhondeh et al., 2019). Thus, it becomes imperative to lower lipid levels in individuals with hyperlipidemia.

Several lipid-lowering drugs in hyperlipidemia therapy include statins and fibrates (Ali et al., 2021). Regrettably, synthetic drugs designed to combat hyperlipidemia and atherosclerosis may entail multiple adverse effects (Ali et al., 2021; Auer et al., 2016). Phytotherapy is a safe solution; herbal medicine may be a suitable alternative to conventional medicine for treating several disorders, including hyperlipidemia and cardiovascular disease (Liu et al., 2011). Additionally, more solutions and products must be found and developed in addition to conventional medications to control lipid levels and prevent cardiovascular disease (Ghatak and Panchal, 2012).

Ginger, scientifically known as *Zingiber officinale* Roscoe, was identified by English botanist William Roscoe and belongs to the Zingiberaceae family. Native to South Asia, this plant possesses perennial rhizomes and thrives in subtropical and tropical regions, scaling altitudes up to 1500 meters above sea level (Enstitüsü, 2015). *Z. officinale* has been a globally utilized spice and herb, notably in India and China, the primary ginger producers. It assumes a crucial role in traditional Chinese herbal medicine and is believed to have spread to other continents during the Roman and Greek epochs (Ali et al., 2008). *Zingiber officinale* has also been extensively applied to treating diverse diseases, including indigestion, nausea, asthma, and muscle pain (Li et al., 2012).

*Citrus limon* L., or lemon, is a plant species belonging to the Rutaceae family that grows in sub-tropical and Mediterranean climates. The *Citrus limon* tree produces a fruit (lemon) widely used for its juice, mainly as a condiment (Kehal, 2013). The juice of this fruit has an acidic flavor and a low pH (2-3) due to its richness in organic acids (Himed and Merniz, 2016). Numerous pharmacological effects of *Citrus limon* have been demonstrated, including antioxidant [12], anti-inflammatory (Amorim et al., 2016), hepatoprotective (Jaiswal et al., 2015), diabetes prevention and anti-obesity (Kim and Ko, 2017), and lipolytic and cholesterol-lowering (Farnier et al., 2021) effects.

This study investigates the anti-hyperlipidemic impact of *Zingiber officinale* and *Citrus limon* juices, individually and in combination, on triton-induced hyperlipidemic mice.

Indeed, while significant strides have been made in understanding the therapeutic potential of natural compounds in the context of hyperlipidemia, there exists a conspicuous gap in the literature concerning the combined effect of *Zingiber officinale* and *Citrus limon* juice extracts on Triton WR-1339-induced acute hyperlipidemia. Our study aimed to bridge this gap by investigating the synergistic impact of these two botanical extracts in a formulated treatment. Furthermore, a comprehensive phytochemical analysis was conducted to identify and characterize the bioactive compounds present in the extracts. Recognizing the pivotal role of HMG-CoA reductase in cholesterol metabolism, a molecular docking study was undertaken to elucidate the potential interactions between the bioactive compounds and the enzyme. Our research sheds light on the therapeutic efficacy of the formulated juice extract and contributes valuable insights into the molecular mechanisms underlying their effects.

## 2. Materials and Methods

### 2.1. Chemicals

The following reagents were purchased from Sigma Chemical Co. (Taufkirchen, Germany): Triton WR-1339, Folin-Ciocalteu, Sodium Carbonate (Na_2_CO_3_), Gallic Acid, Sodium Nitrate (NaNO_3_), Aluminum Chloride (AlCl_3_), Sodium Hydroxide (NaOH), Quercetin, Sodium Phosphate (Na_3_PO_4_).

### 2.2. *Zingiber officinale* Roscoe (Ginger)

*Zingiber officinale* Roscoe (Ginger) rhizomes at the mature stage were purchased from an herbalist (Oujda, Morocco) and well-cleaned with distilled water to remove any dirt. Professor Fennane Mohammed, a qualified botanist from the Scientific Institute of Rabat (Morocco), identified the plant material. A reference voucher specimen with the reference number was kept in the herbarium of Mohamed First University’s Faculty of Sciences in Oujda, Morocco (HUMPOM-352).

### 2.3. *Citrus limon* L. (Lemon)

At the mature stage, the lemon fruits (*Citrus limon* L.) were purchased from a local market (Oujda, Morocco), washed adequately with distilled water, and authenticated by Professor Fennane Mohammed. A reference voucher specimen has been deposited in the herbarium of the Faculty of Sciences of Mohamed First University (Oujda, Morocco) under the reference number (HUMPOM-450).

### 2.4. Zingiber officinale and Citrus limon juice extraction

#### 2.4.1. *Zingiber officinale* juice extraction

To make ginger juice (GJ), 500 g of ginger rhizomes were cut into smaller pieces of approximately 1 cm and ground in a blender at 25 °C. The resulting “GJ” was filtered through “Whatman Grade 40 high-purity” filters (medium nominal particle retention rating of 8 µm) and concentrated using a rotating vacuum evaporator (Heidolph Rotary Evaporator, Laborota 4010 intensive condenser G4) before being stored at - 20 °C until its use. An extraction yield of 2.3% (w/w) was obtained.

#### 2.4.2. *Citrus limon* juice extraction

Lemons were washed and cut into small pieces weighing 500 g before being ground for 1-2 minutes using a blender to extract the lemon juice (LJ). Then, the same procedures were performed precisely as those in ginger juice extraction, and a yield of 2 % (w/w) was attained.

#### 2.4.3. Formulation preparation

The formulation was prepared by combining 50% *Z. officinale* juice (GJ) with 50% *C. limon* juice (LJ) and then used to prepare any concentration or dose needed.

### 2.5. Phytochemical Analysis

The phytochemical properties of *Z. officinale* and *C. limon* were analyzed to identify lead compounds and components implicated in observed physiological effects. The distinctive biological activities of these plants can be determined through their phytochemical properties.

The phytochemical studies done in this work were: “The total phenolic quantification”, “the total flavonoids determination”, and “the HPLC analysis”.

#### 2.5.1. Total Phenolics Quantification

The quantification of total phenolics in *Z. officinale* and *C. limon* juices was carried out using a slightly modified method as described by Ainsworth & Gillespie (2007), and the calculation of the amount of the polyphenol was based on a calibration curve of the gallic acid. Dilutions of each juice (500 µL) were combined with 2,500 µL of Folin-Ciocalteu reagent (0.2 N) and 2,000 µL of sodium carbonate (Na_2_CO_3_) solution (7.5% w/v) at 25 °C. After 15 minutes of incubation in the dark, absorbance was measured at 765 nm. The process for the blank was the same, except that solvent was used instead of juice. Total phenolic content was expressed in micrograms of gallic acid equivalents per milligram of extract (mg GAE/mg extract) using a calibration curve prepared with gallic acid.

#### 2.5.2. Total Flavonoids Determination

The determination of flavonoids in *Z. officinale* and *C. limon* juices was conducted following the method outlined by (Chen et al., 2015), and the calculation of the flavonoid amount was based on a calibration curve of the quercetin. Dilutions of each juice (200 µL) were mixed with 1 mL of distilled water and 50 µL of sodium nitrate (NaNO_3_) solution (5% w/v). The mixture was then combined and left to incubate at room temperature for 5 minutes. Subsequently, 120 µL of aluminum chloride (AlCl_3_) solution (10% w/v) was added. After another 5 minutes of incubation at 25 °C, 400 µL of sodium hydroxide (NaOH) solution (1M) was introduced to the reaction, followed by absorbance measurement at 430 nm. The procedure for preparing the blank was identical, except that solvent was used instead of juice. The total flavonoid content was quantified in micrograms of quercetin equivalents per milligram extract (mg E.Q./mg).

#### 2.5.3. High-Performance Liquid Chromatography (HPLC)

High-Performance Liquid Chromatography (HPLC) was employed to qualitatively evaluate the phenolic compounds in *Z. officinale* and *C. limon* juices. The analysis utilized a Waters Alliance 2695 HPLC system and a range of standards, including 4-gingerol, 6-gingerol, 6-gingediol, eriodictyol, rutin, hesperidin, and isorhamnetin.

In the previous study (17), HPLC analysis of both juices was carried out. Chromatographic separation was achieved using a reversed-phase C18 column (250 × 4.6 mm, five μm pore size). The mobile phase comprised two solvents: solvent A (a mixture of water and formic acid in a 90:10 v/v ratio) and solvent B (a mixture of water, methanol, and acetonitrile in a 40:50:10 v/v ratio). The eluent flow rate was kept at 1 mL/min, and the injection volume was set at 20 μL. All analyses were conducted at room temperature.

Standard and extract solutions of *Z. officinale* and *C. limon* (GJ, LJ, and F) were dissolved in methanol and filtered through a 0.45 μm Millipore membrane.

### 2.6. Antioxidant study of *Z. officinale* and *C. limon* Juices

#### 2.6.1. Phosphomolybdenum assay

The total antioxidant capacity (TAC) was evaluated using the phosphomolybdenum method, following the previously described by El Moussaoui et al. (2019) was meticulously adopted to assess the extract total antioxidant capacity (TAC), employing a well-established protocol. Specifically, 0.1 mL of the extract was combined with a reagent mixture containing 0.6 M sulfuric acid, 28 mM sodium phosphate, and 4 mM ammonium molybdate in a reaction tube. The tubes were securely sealed and incubated in a water bath at 95°C for 90 minutes to facilitate the reaction. Following incubation, the absorbance of the solutions was measured at 695 nm against a reagent blank. The results were quantified as milligrams of ascorbic acid equivalents per gram of dry weight (mg AAE g⁻¹ DW), providing a standardized measure of the extract’s antioxidant capacity.

#### 2.6.2. ABTS assay

The radical-scavenging activity of ABTS [2,2′-azino-bis(3-ethylbenzothiazoline-6-sulfonic acid)] was determined using the method established by Re et al. (1999), with slight modifications. Briefly, an aqueous solution of potassium persulfate (2.45 mM) was mixed with a 7 mM ABTS aqueous solution to prepare the assay reagent, which was then left in the dark at room temperature overnight to allow radical generation. Before analysis, the reagent was diluted with ethanol (EtOH) to obtain an absorbance of 0.70 (± 0.01) at 734 nm. For the assay, 100 µL of the extract at varying concentrations was added to 2 mL of the ABTS solution. The mixture was incubated for 10 minutes, and absorbance was measured at 734 nm.

Ethanol served as the control, while calibration was performed using a standard curve of ascorbic acid solutions. The percentage inhibition was calculated using the formula:

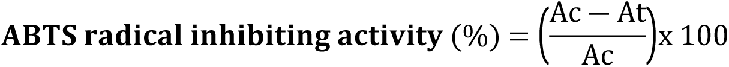

Where “Ac” represents the absorbance of the control (without the extract) and “At” represents the absorbance of the test sample.

Results were expressed as IC_50_ values, indicating the sample concentration required to inhibit 50% of the ABTS radicals.

#### 2.6.3. DPPH radical scavenging assay

Thanks to its stability as a free radical and the simplicity of its analysis, DPPH (2,2-diphenyl-1-picrylhydrazyl) has become a popular tool for quickly and efficiently evaluating antioxidant activity (McCune and Johns, 2002). Subhashini et al. (2011) detailed the experimental procedure to explore DPPH’s radical scavenging ability. Specifically, 1 mL of the plant’s etheric extract was mixed with 1 mL of a DPPH radical solution, achieving a final DPPH concentration of 0.025 g/L. After thorough stirring, the mixture was left to rest for 30 minutes, and its absorbance was then recorded at 517 nm. Ascorbic acid served as the standard reference. The percentage of DPPH radical inhibition was calculated using an equation, as outlined by Manzocco et al. (1998) (Manzocco et al., 1998)

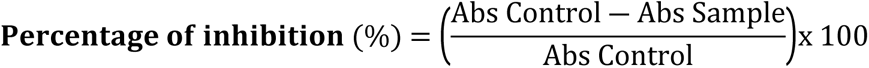

### 2.7. Evaluation of the Protective Effect of GJ, LJ, and BHA Against Plasma Lipoprotein Oxidation

The protective effects of ginger juice (GJ), lemon juice (LJ), and butylated hydroxyanisole (BHA) against plasma lipoprotein oxidation were assessed by quantifying malondialdehydes (MDA), secondary products of lipoprotein oxidation, as thiobarbituric acid reactive substances (TBARS), following the method described by Bekkouch et al. (2019) (Bekkouch et al., 2019).

Lipoprotein-rich plasma, used as the oxidation substrate, was collected from mice treated with 400 mg/kg of Triton WR-1339 for 10 hours. As previously determined, this plasma contained approximately 100 ± 5 mg/dL of LDL cholesterol.

The oxidative process was initiated using copper sulfate (CuSO₄) according to the following protocol:

- **Control group**: 40 µL of lipoprotein-rich plasma incubated with distilled water.
- **Oxidized lipoproteins group**: 40 µL of lipoprotein-rich plasma incubated with 10 µL of CuSO₄ (0.3 mg/mL).
- **GJ-treated lipoproteins group**: 40 µL of lipoprotein-rich plasma incubated with 10 µL of CuSO₄ solution and *Zingiber officinale* juice at concentrations of 5, 12.5, 25, 50, 100, and 200 µg/mL.
- **LJ-treated lipoproteins group**: 40 µL of lipoprotein-rich plasma incubated with 10 µL of CuSO₄ solution and *Citrus limon* juice at concentrations of 5, 12.5, 25, 50, 100, and 200 µg/mL.
- **Formulation-treated lipoproteins group**: 40 µL of lipoprotein-rich plasma incubated with 10 µL of CuSO₄ solution and ginger and lemon Formulation at concentrations of 5, 12.5, 25, 50, 100, and 200 µg/mL.
- **BHA-treated lipoproteins**: 40 µL of lipoprotein-rich plasma incubated with 10 µL of CuSO₄ solution and BHA at concentrations of 5, 12.5, 25, 50, 100, and 200 µg/mL.

After vigorous mixing, the preparations were incubated at 37°C for 24 hours. Subsequently, 500 µL of 20% trichloroacetic acid and 500 µL of 0.8% thiobarbituric acid were added to each sample. The reaction mixtures were heated at 95°C for 30 minutes, then cooled to room temperature. The absorbance was measured at 532 nm.

TBARS levels were calculated and expressed as MDA equivalents, determined from a calibration curve. All measurements were performed in triplicate to ensure accuracy and reliability.

### 2.8. Hypolipidemic Effect of *Z. officinale* and *C. limon* Juices in Mice Model

#### 2.8.1. Animals

90 adult male albino mice weighing 25–30 g were raised in the biology department’s animal house at the Faculty of Sciences in Oujda following international guidelines established by the National Institutes of Health of the United States (NIH Publication No. 85–23, revised 1985) for the protection and usage of laboratory animals (20). The Faculty of Sciences Institutional Review Board of Oujda University, Morocco, approved the study (03/21-LBBEH-15 and March 5, 2021).

The animals were maintained in a temperature-controlled room (22 ± 2°C) with a 12 h/12 h light/dark cycle and provided food and water ad libitum.

#### 2.8.2. Induction of Hyperlipidemia by Triton WR-1339

Triton WR-1339 is a nonionic detergent widely used to induce acute hyperlipidemia in animal models to study cholesterol and triglyceride metabolism (Ghatak and Panchal, 2012). The accumulation of plasma lipids caused by this detergent is due to inhibiting lipoprotein lipase activity (Scharwey et al., 2013). Animals were injected with a dose of 200 mg/kg, according to (Schurr et al., 1972).

Adult male *Albino* mice were divided into nine groups of 10 animals each:

- **Group 1:** normolipidemic control animals gavaged with distilled water (NCG).
- **Group 2:** animals receiving Triton WR-1339 (200 mg/kg in 9‰ NaCl; pH= 7.4) by intraperitoneal injection and gavaged with distilled water (HCG).
- **Group 3:** Animals injected with Triton WR-1339 (200 mg/kg) and administered with *Z. officinale* juice (250 mg/kg) (GJTG 1).
- **Group 4:** Animals injected with Triton WR-1339 (200 mg/kg) and treated with *Z. officinale* juice (500 mg/kg) (GJTG 2).
- **Group 5:** Animals injected with Triton WR-1339 (200 mg/kg) and treated with *C. limon* juice extract (250 mg/kg) (LJTG 1).
- **Group 6:** Animals are given Triton WR-1339 (200 mg/kg) and administered with *C. limon* juice extract (500 mg/kg) (LJTG 2).
- **Group 7:** Animals receiving Triton WR-1339 (200 mg/kg) and treated with the formulation of the *Z. officinale* and the *C. limon* juices (250 mg/kg) (FTG 1).
- **Group 8:** Animals fed Triton WR-1339 (200 mg/kg) and gavaged with the formulation from *Z. officinale* juice and *C. limon* juices (500 mg/kg) (FTG 2).
- **Group 9:** animals receiving Triton WR-1339 (200 mg/kg) and gavaged with Atorvastatin (10 mg/kg) (ATG).

#### 2.8.3. Biochemical Analysis

##### a) Blood Sampling

Following 24 hours post-injection, mice were gently anesthetized, and then blood was collected from the retroorbital plexus using heparinized Eppendorf tubes, subsequently undergoing centrifugation at 1500 rpm for 15 minutes. This process led to the isolation of plasma, which was then transferred into separate Eppendorf tubes for lipid quantification.

##### b) Determination of Total Cholesterol (TC)

The enzymatically colorimetric approach was used to measure the plasma levels of total cholesterol (TC) using a commercially available kit (Biosystems Kit, Barcelona, Spain, REF: 12505). For instance, 1 mL of the enzymatic total cholesterol reagent was mixed with 10 µL of plasma samples for 10 min at 37°C. The absorbance was then measured at 510 nm, and the TC was determined according to the manufacturer’s instructions.

##### c) Determination of Triglycerides (TG)

Plasma triglycerides were determined using a commercial kit (Biosystems Kit, Barcelona, Spain, REF: 12558). The enzymatic triglyceride reagent was mixed with 10 µL of plasma samples for 10 min at 37°C. The absorbance was then determined at 510 nm, and the TG was determined according to the manufacturer’s instructions.

##### d) High-Density Lipoprotein Cholesterol (HDLc) and Low-Density Lipoprotein Cholesterol (LDLc) Assay

The HDL cholesterol fraction was measured after LDL cholesterol and VLDL lipoproteins were precipitated by phosphotungstic acid (PTA) in the presence of magnesium chloride (MgCl_2_). In a tube, 20 µL of plasma and 10 µL of the “PTA/MgCl_2_ “reagent are combined, and the mixture is then centrifuged at 5,000 rpm for 15 minutes after standing for 10 minutes. The same procedure was used for the total cholesterol assay, which measures HDL cholesterol in the supernatant. Blood levels of low-density lipoprotein cholesterol (LDLc) were determined using the Friedewald formula. The same technique was employed to quantify HDLc.

### 2.9. Acute toxicity study in mice

The acute oral toxicity study was conducted per the Organization for Economic Cooperation and Development (OECD) guidelines 423 (OECD Guideline, 2001).

96 mice were used, divided into 16 groups of 6 mice each (3 males and three females per group). The first group served as the control and received only distilled water. The remaining 15 groups were administered increasing doses of the GJ, LJ, and the formulation F at 2, 4, 6, 8, and 10 g/kg of body weight.

Following the oral administration of the GJ, LJ, and formulation F, the mice were carefully observed individually for the first 30 minutes. Regular monitoring continued over the next 24 hours, with special attention given to the critical first 4 hours. This observation period was maintained daily for 14 days to assess potential toxic effects thoroughly.

### 2.10. Subacute Toxicity Study

#### 2.10.1. Animal protocol

Eleven groups of rats were created to assess subacute toxicity, each with six individuals (three males and three females). The groups were organized as follows:

a. Control Group: Received distilled water orally for 30 days.
b. Test Groups: Administered different doses of each extract (500, 1000, or 2000 mg/kg body weight) by gavage daily for 30 days.

The dose levels were carefully determined based on the LD_50_ and guidelines provided in the Organisation for Economic Cooperation and Development (OECD) document 407.

Throughout the 30 days, the animals were monitored daily to check for any changes in their general health or signs of toxicity. Body weight measurements were taken on days 0, 7, 14, 21, and 28 to track fluctuations.

At the end of the study, the animals fasted overnight before blood samples were collected under anesthesia from the abdominal aorta. Blood was drawn into two types of tubes:

`o` *EDTA-coated tubes: These were used immediately for hematological analysis*.
`o` Plain tubes: These were centrifuged at 3000 rpm at 4°C for 10 minutes to separate the serum, which was later used for biochemical testing.

#### 2.10.2. Serum Biochemistry

To gain a deeper understanding of the physiological impacts of the treatment, we analyzed serum samples using an automated chemistry analyzer (COBAS INTEGRA^®^ 400 Plus). This process allowed us to measure various important biochemical markers that reflect the health of key organs. Albumin (ALB) levels were assessed as a general indicator of liver function and nutritional status. Alkaline phosphatase (ALP) was analyzed to check for potential liver or bone health issues. Furthermore, we measured alanine aminotransferase (ALT) and aspartate transaminase (AST), two enzymes commonly used to detect liver stress or damage.

We also evaluated bilirubin (BIL), which can signal liver function abnormalities or hemolytic activity. To assess lipid metabolism, cholesterol (CHOL) and triglycerides (TRGL) were measured, providing insights into the subjects’ cardiovascular and metabolic health. Finally, creatinine (CRE) and urea (URE) were analyzed to gauge kidney function and overall protein metabolism.

Combining all these parameters gave us a detailed picture of how the treatment influenced the animals’ internal systems. This comprehensive approach helped us identify potential toxic effects, particularly on the liver, kidneys, and metabolic functions, ensuring a thorough evaluation of the treatment’s safety.

#### 2.10.3. Hematologic Markers

The blood analysis was carried out using an automated hematology analyzer (Abacus 380 Hematology Analyzer) to provide a detailed profile of the blood composition. The examination focused on several key indicators of blood health and function.

White blood cell (WBC) counts were measured to evaluate immune system activity, alongside red blood cell (RBC) counts, which provide insight into oxygen transport capacity. Hemoglobin (HGB) and hematocrit (HCT) levels were analyzed to evaluate oxygen-carrying capacity and overall blood volume.

To further explore red blood cell characteristics, the mean corpuscular volume (MCV), mean corpuscular hemoglobin (MCH) and mean corpuscular hemoglobin concentration (MCHC) were measured. This provided details on cell size, hemoglobin content, and variability.

Platelet (PLT) counts and mean platelet volume (MPV) were also recorded, offering insights into clotting function and platelet activity.

This thorough analysis offered a comprehensive overview of the blood’s cellular components, ensuring a detailed understanding of the treatment’s effects on hematological health.

### 2.11. Molecular Docking Study

After having shown the beneficial effect of *Z. officinale* and *C. limon* juices on cholesterol levels, and to deepen our knowledge of the signaling pathway involved in this effect, we carried out an in silico study which consisted in determining the affinity between the most abundant molecules in the two extracts with the enzyme responsible for cholesterol synthesis : 3-hydroxy-3-methyl-glutaryl-coenzyme A reductase (HMG-CoA-reductase) which was downloaded from the “PDB” database known as “1DQ9” 3D in SDF format The grid box had a grid card size of 40 × 40 × 40 grid points centered at 14.18 × 02.51 × 11.49, for drug development, molecular clustering has been used as an excellent tool for predicting the binding modes between ligand and receptor, The molecules were downloaded from PubChem and were prepared using “PyMOL” (www.pymol.org) and also AutoDock Vina” (www.vina.scripps.edu), the choice of molecules was made according to the content of these derivatives in the two extracts according to an HPLC study previously done in a previous work (Bekkouch et al., 2022), and based on this work, for the *C. limon* juice we have chosen: eriodictyol, rutin, isorhamnetin, hesperidin and for the ginger extract we have chosen: 6-gingerol, 6-gingediol, 4-gingerol, in addition to the reference molecule, which is Atorvastatin.

The binding between the enzyme and the molecules occurred thanks to the auto-dock software, which allowed us to determine the affinity between the enzyme and each molecule. Then, the critical points were visualized in 2D and 3D thanks to the Biovia Discovery Studio software.

### 2.12. Statistical Analysis

The Student’s t-test was used to analyze the collected experimental data. GraphPad Prism 9.0.0 (GraphPad Prism Software, Inc.: San Diego, CA, USA) was utilized using an unpaired Student’s t-test for statistical significance between the two groups. Then, the analysis of variance (ANOVA) was followed by Tukey’s pairwise comparison test at a 95% confidence interval (p < 0.05). The P values were considered statistically significant because they were lower than 0.05. The results are shown as mean ± SEM.

## Results

### 3.1. Determination of Total Phenolic Content

The determination of the total phenolic content by the method of Folin-Ciocalteu revealed the presence of 18.48 ± 1.14 mg and 25.23 ± 1.54 mg equivalent of gallic acid equivalent/g DW in the juices of *Z. officinale* juice (GJ) and *C. limon* juice (LJ) respectively (table 1), the calculation of these quantities was due to a calibration curve of the gallic acid (figure 1).

**Figure 1.**
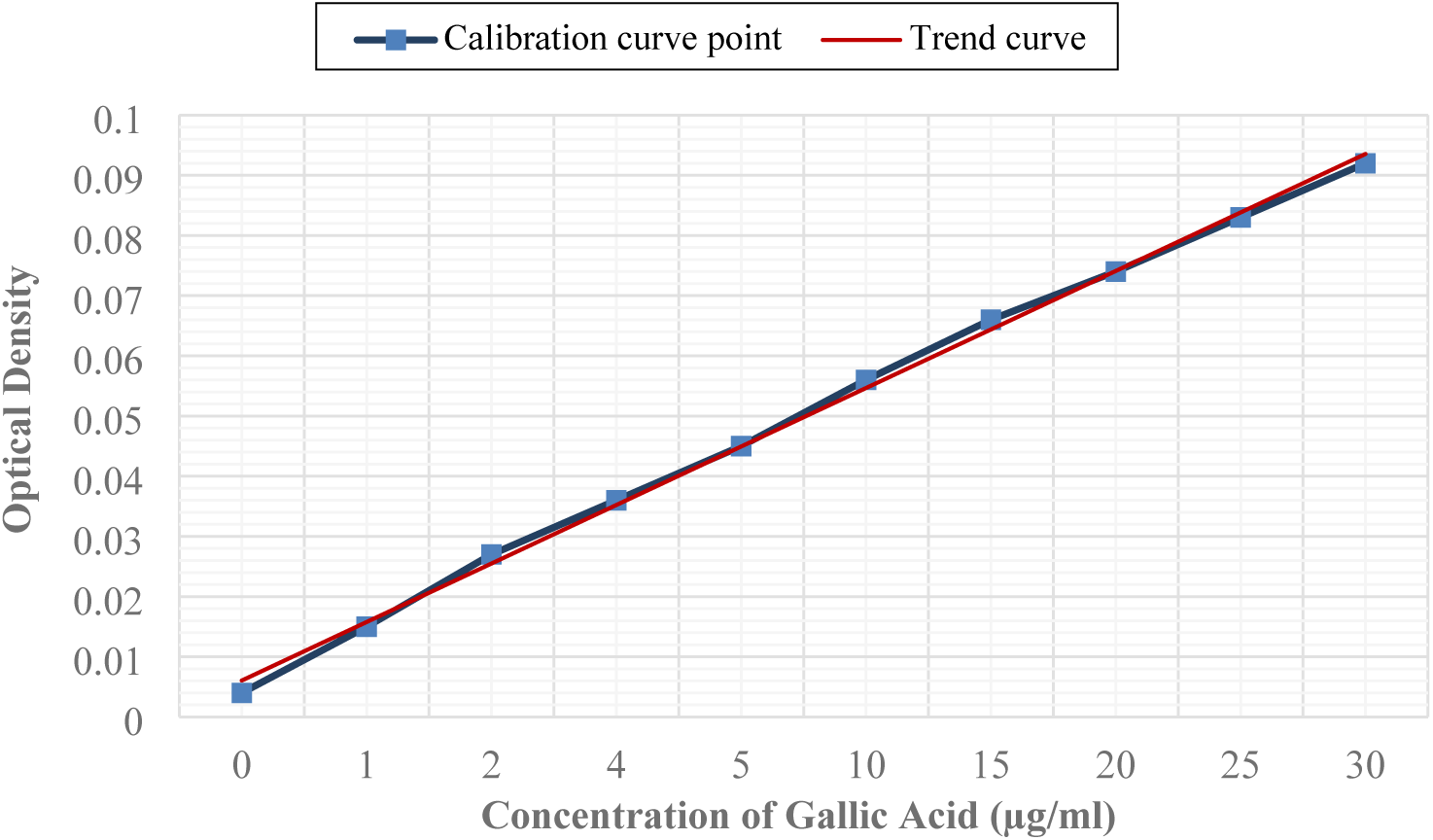
Calibration curve of Gallic Acid.

**Table 1.**
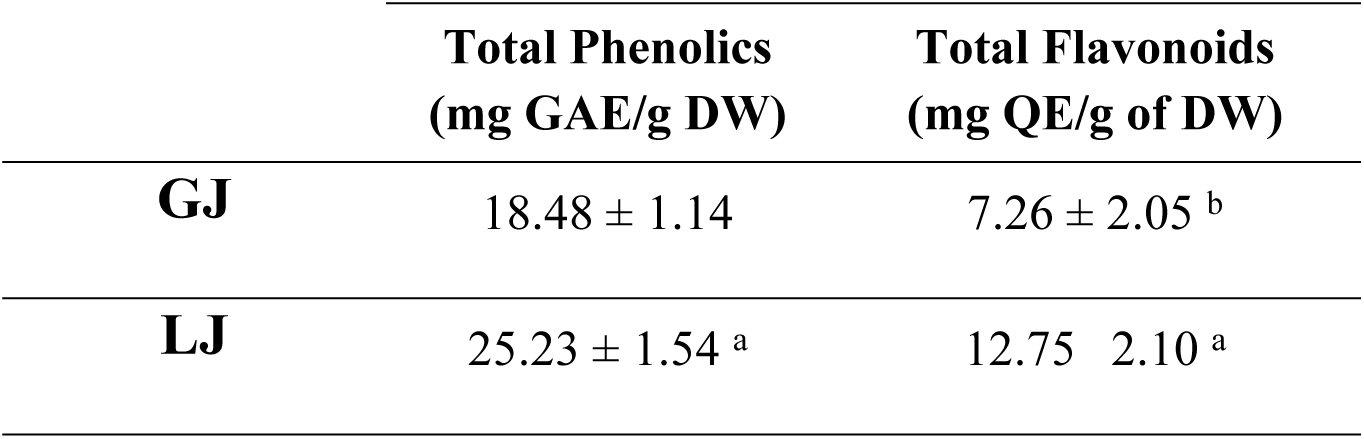
Total phenolics and flavonoids quantification of ginger juice (GJ) and lemon juice (LJ).

|  | Total Phenolics<br>(mg GAE/g DW) | Total Flavonoids<br>(mg QE/g of DW) |
| --- | --- | --- |
| **GJ** | $18.48 \pm 1.14$ | $7.26 \pm 2.05^b$ |
| **LJ** | $25.23 \pm 1.54^a$ | $12.75 \pm 2.10^a$ |

### 3.2. Determination of Total Flavonoids Content

The total flavonoid content revealed that “GJ” and “LJ” contain 7.26 ± 2.05 and 12.75 ± 2.10 mg equivalent of quercetin/g, respectively (Table 1). These quantities were calculated due to a calibration curve of the gallic acid (figure 2).

**Figure 2.**
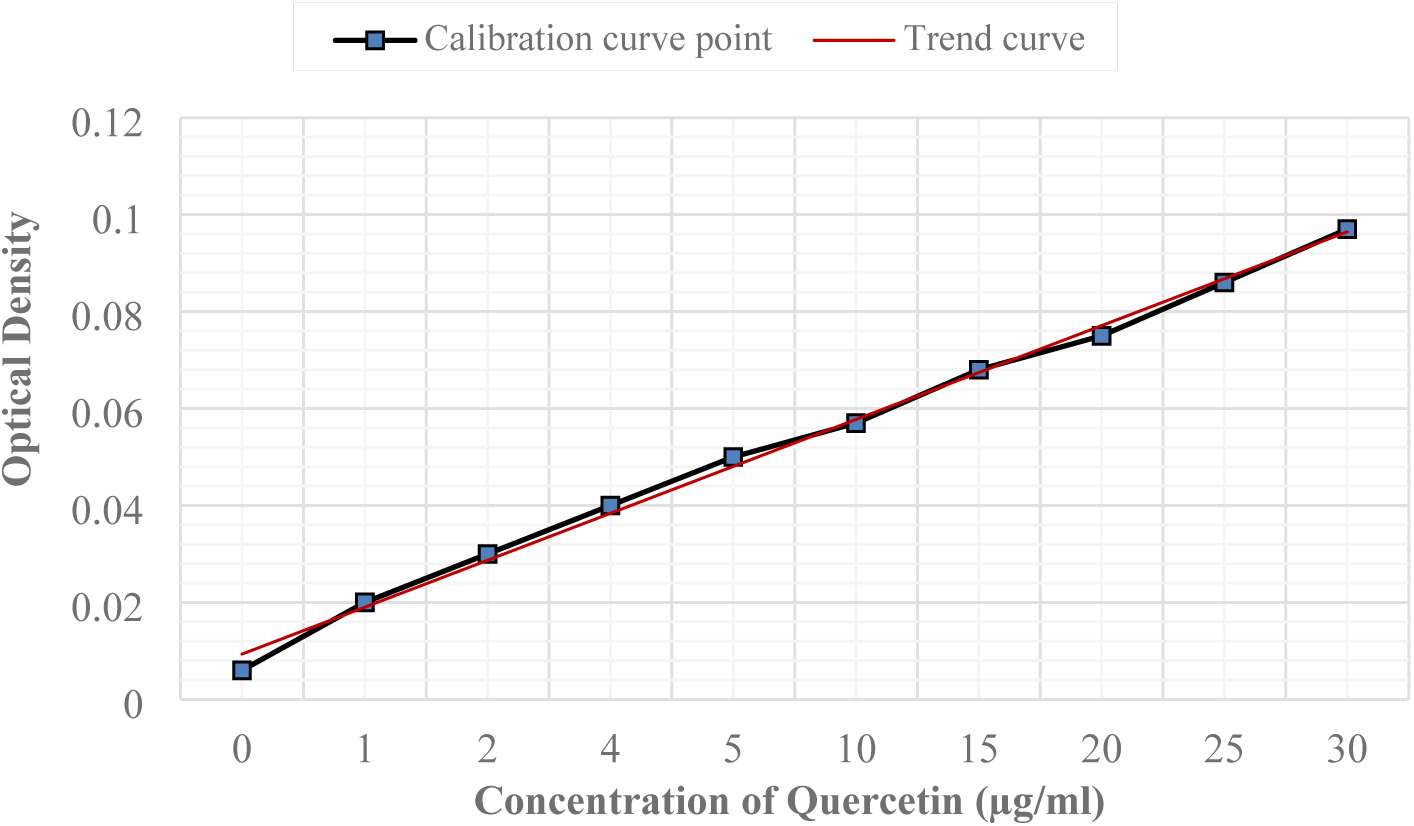
Calibration curve of quercetin.

Columns with similar alphabets (letters a or b) are not significantly different (p < 0.05). Values are expressed as means ± SEM from triplicates. mg GAE/g DW: milligram equivalent of gallic acid equivalent per gram of dry weight, mg QE/g of DW: milligram equivalent of quercetin equivalent per dry weight.

### 3.3. HPLC Chromatography of *Zingiber officinale* and *Citrus limon* Juices

The results of the analysis of Z. officinale (GJ) and C. limon (LJ) juices using High-Performance Liquid Chromatography (HPLC) revealed distinct profiles for each.

In the case of “GJ,” the main compounds identified were 6-gingerol, 4-gingerol, and 6-gingediol (**Figure 3-A**, **Table 2-a**).

**Figure 3.**
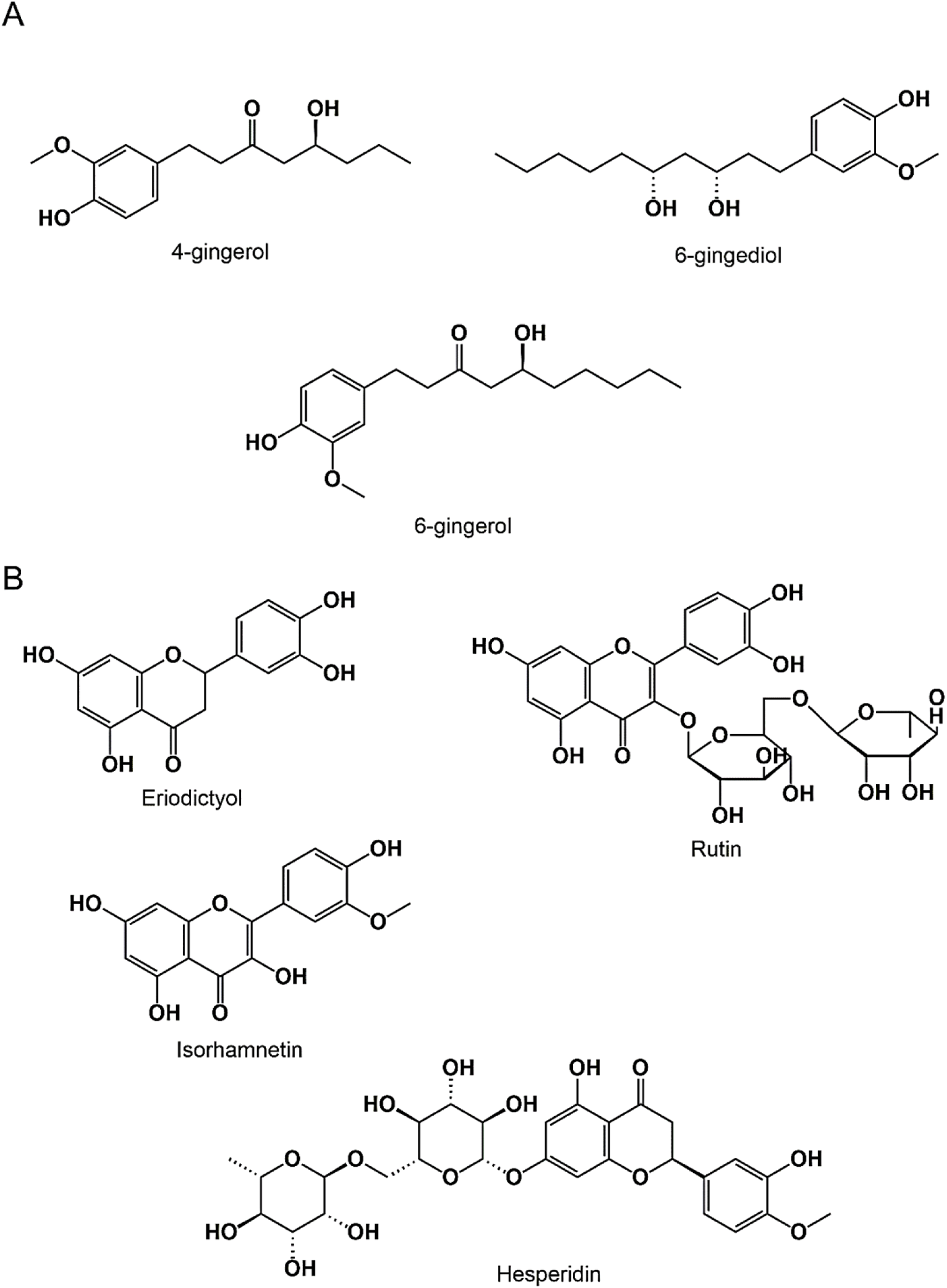
Chemical structure of *Zingiber officinale* (A) and *Citrus limon* (B) phenolic compounds (drawn using “ChemDraw 12.0”)

**Table 2.**
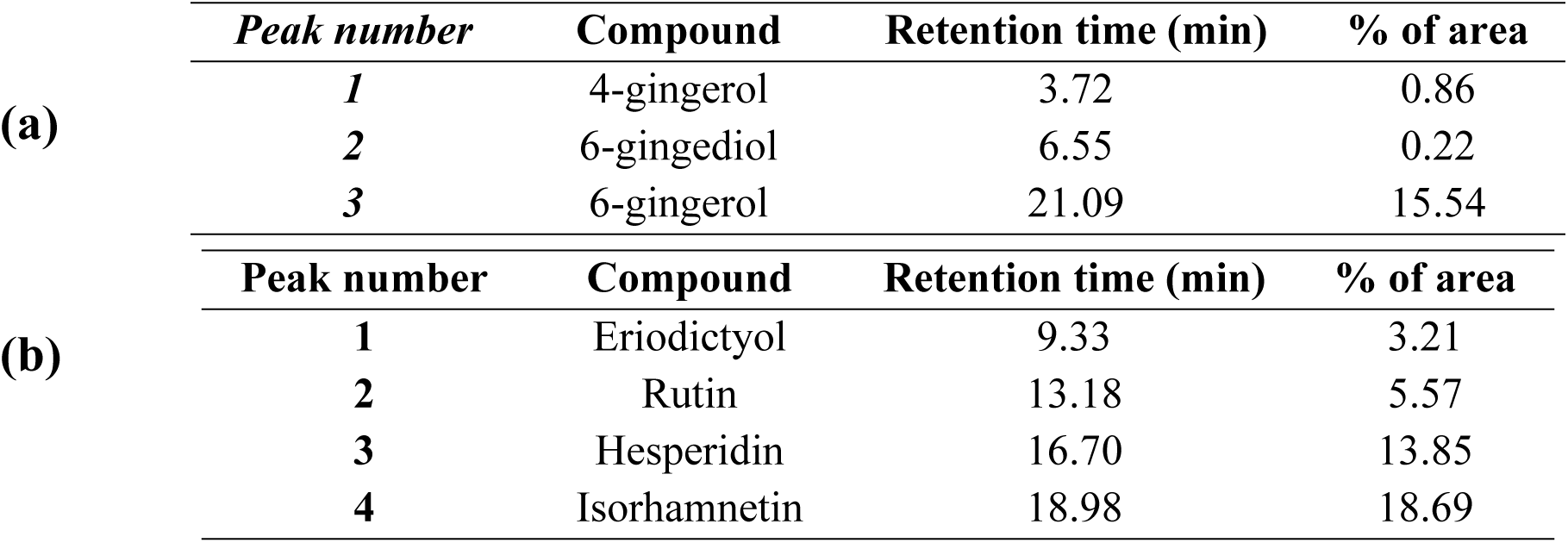
Chemical composition of *Zingiber officinale* juice (a) and *Citrus limon* (b)

The HPLC analysis of “LJ” showed the existence of several molecules, and this analysis allowed us to prove the existence of hesperidin, rutin, isorhamnetin, and eriodictyol (Figure 3-B, Figure 4-A, Table **3**).

**Figure 4.**
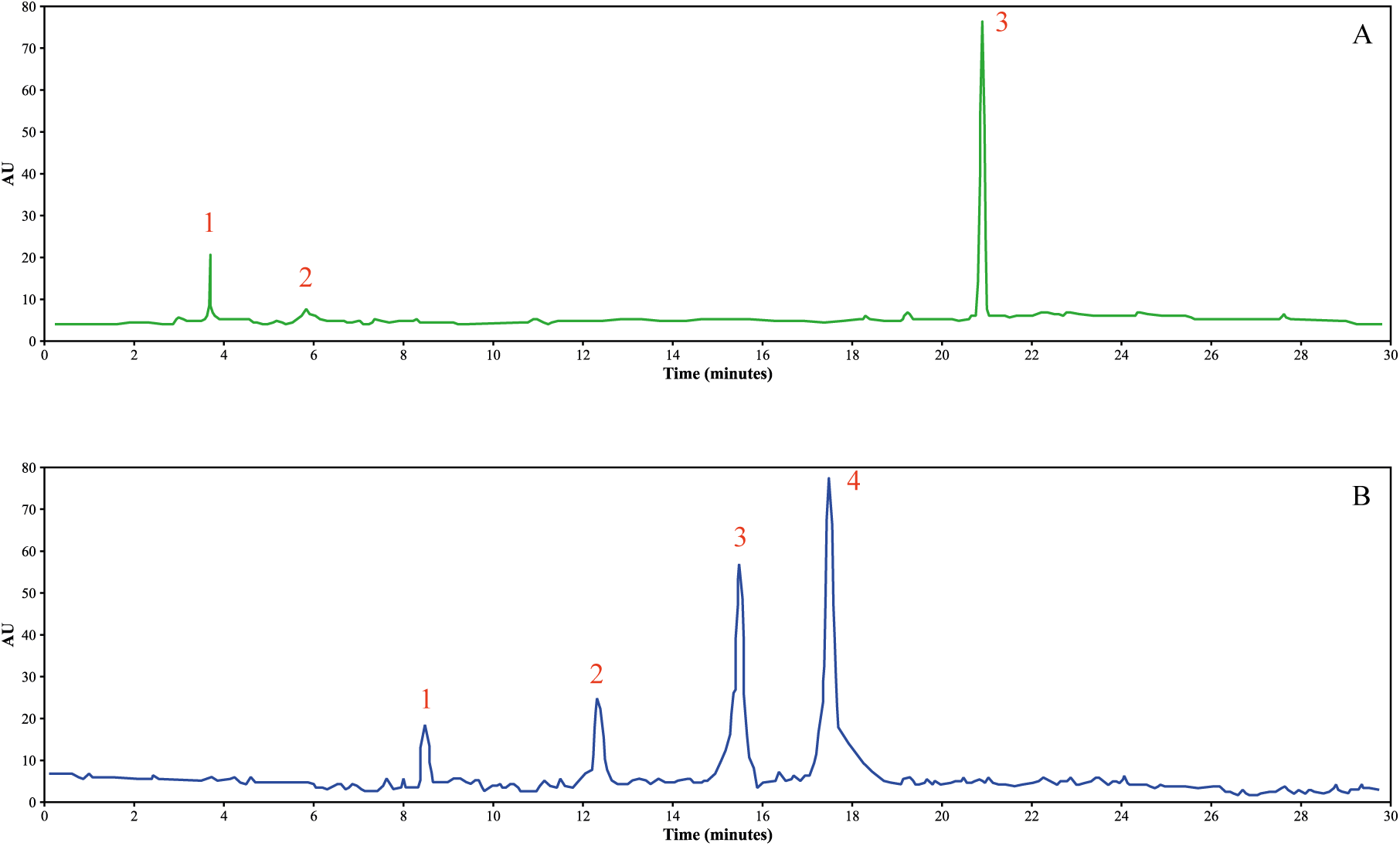
HPLC chromatographic profiles of *Zingiber officinale* (A) and *Citrus limon* (B) juices*.

**Table 3.** IC_50_ Values of Antioxidant Activities of *Zingiber officinale* and *Citrus limon* Extracts and Their Formulation Across DPPH, ABTS, TAC, and FRAP Assays.

|  | **Ginger juice (GJ)** | **Lemon juice (LJ)** | **Formulation (F)** | **Ascorbic acid** |
| --- | --- | --- | --- | --- |
| **DPPH** | 38.91 ± 1.47 | 43.14 ± 1.58 | 44.94 ± 1.02 | 40.11 ± 0.76 |
| **ABTS** | 31.78 ± 0.63 | 33.1 ± 1.74 | 30.92 ± 2.0 | 34.27 ± 1.39 |
| **TAC** | 58.43 ± 2.2 | 59.49 ± 2.34 | 63.9 ± 1.11 | 60.14 ± 0.77 |
| **FRAP** | 74.59 ± 1.85 | 76.91 ± 2.14 | 77.9 ± 1.76 | 75.12 ± 2.24 |

Data are expressed in (Mean ± SEM)

On the other hand, the HPLC analysis of *C. limon* juice exhibited a more diverse range of compounds. This thorough analysis confirmed the presence of hesperidin, rutin, isorhamnetin, and eriodictyol in the *Citrus limon* juice (**figure 3-B**, **figure 4-B**, **table 2-b**).

### 3.4. Antioxidant study of *Zingiber officinale* and *Citrus limon* juices

Table 3 elucidates the results of the antioxidant activities of *Zingiber officinale* (ginger) *and Citrus limon* (lemon) extracts across four antioxidant assays: DPPH, ABTS, TAC, and FRAP. The data demonstrate the effectiveness of the extracts and their formulation (F) compared to ascorbic acid, a standard antioxidant. Lower IC_50_ values correspond to higher antioxidant activity.

The DPPH radical scavenging assay revealed that ascorbic acid exhibited the most potent activity (IC_50_ = 40.11 ± 0.76 µg/mL), followed closely by the formulation (IC_50_ = 44.94 ± 1.02 µg/mL). Interestingly, lemon juice (IC_50_ = 43.14 ± 1.58 µg/mL) displayed a slightly higher IC_50_ than ginger juice (IC_50_ = 38.91 ± 1.47 µg/mL), suggesting comparable radical scavenging capabilities, though the formulation demonstrated an evident synergistic effect.

Similarly, the ABTS assay reinforced the superior activity of ascorbic acid (IC_50_ = 34.27 ± 1.39 µg/mL). What is particularly noteworthy is that ginger juice (IC_50_ = 31.78 ± 0.63 µg/mL) and lemon juice (IC_50_ = 33.10 ± 1.74 µg/mL) showed significant antioxidant activity, whereas the formulation (IC_50_ = 30.92 ± 2.00 µg/mL) exhibited slightly reduced activity compared to the individual extracts. This observation might be attributed to complex interactions among the bioactive compounds in the juices, which could either enhance or dampen their collective effect.

Moving on to the TAC assay, the pattern remained consistent, with ascorbic acid demonstrating the highest antioxidant capacity (IC_50_ = 60.14 ± 0.77 µg/mL). Remarkably, the formulation (IC_50_ = 63.90 ± 1.11 µg/mL) also displayed robust activity, slightly surpassing lemon juice (IC_50_ = 59.49 ± 2.34 µg/mL) and ginger juice (IC_50_ = 58.43 ± 2.20 µg/mL). These relatively close IC_50_ values underscore the uniform antioxidant potential of the individual components and their combined formulation.

Finally, the FRAP assay provided additional insights into the reducing power of the samples. Not surprisingly, ascorbic acid exhibited the most potent reducing power (IC_50_ = 75.12 ± 2.24 µg/mL). Among the juices, ginger juice (IC_50_ = 74.59 ± 1.85 µg/mL) stood out with the highest reducing power, followed by lemon juice (IC_50_ = 76.91 ± 2.14 µg/mL). However, the formulation (IC_50_ = 77.90 ± 1.76 µg/mL) displayed slightly lower activity, reflecting potential antagonistic interactions affecting its reducing power.

In summary, the results of the four assays collectively confirm that ascorbic acid serves as a robust antioxidant benchmark. While the individual juices of *Zingiber officinale* and *Citrus limon* exhibited notable antioxidant activities, their formulation demonstrated significant, albeit not consistently superior, activity. This variability could be explained by interactions among the bioactive compounds in the formulation, which might modulate antioxidant efficacy. Nevertheless, the formulation exhibits strong antioxidant potential, underscoring its applicability in nutraceutical and functional food products. Therefore, further investigations are warranted to better understand the mechanisms of interaction and optimize the antioxidant properties of such combinations.

**Figure 5** presents the antioxidant effects of *Zingiber officinale* juice (GJ), *Citrus limon* juice (LJ), their combined formulation (F), and butylated hydroxyanisole (BHA) as a positive control on lipid-rich plasma (Ox-LRP) subjected to oxidative stress induced by Triton WR-1339 injection. The extent of lipid peroxidation was assessed using the TBARS assay, with malondialdehyde (MDA) levels measured as an indicator of lipid peroxidation. The control group, representing baseline conditions without oxidative stress, exhibited low MDA levels, confirming the stability of lipid structures under normal physiological conditions. In contrast, the Ox-LRP group displayed consistently elevated MDA levels at all tested concentrations, highlighting the substantial lipid peroxidation induced by Triton WR-1339 and validating the experimental oxidative stress model.

**Figure 5.**
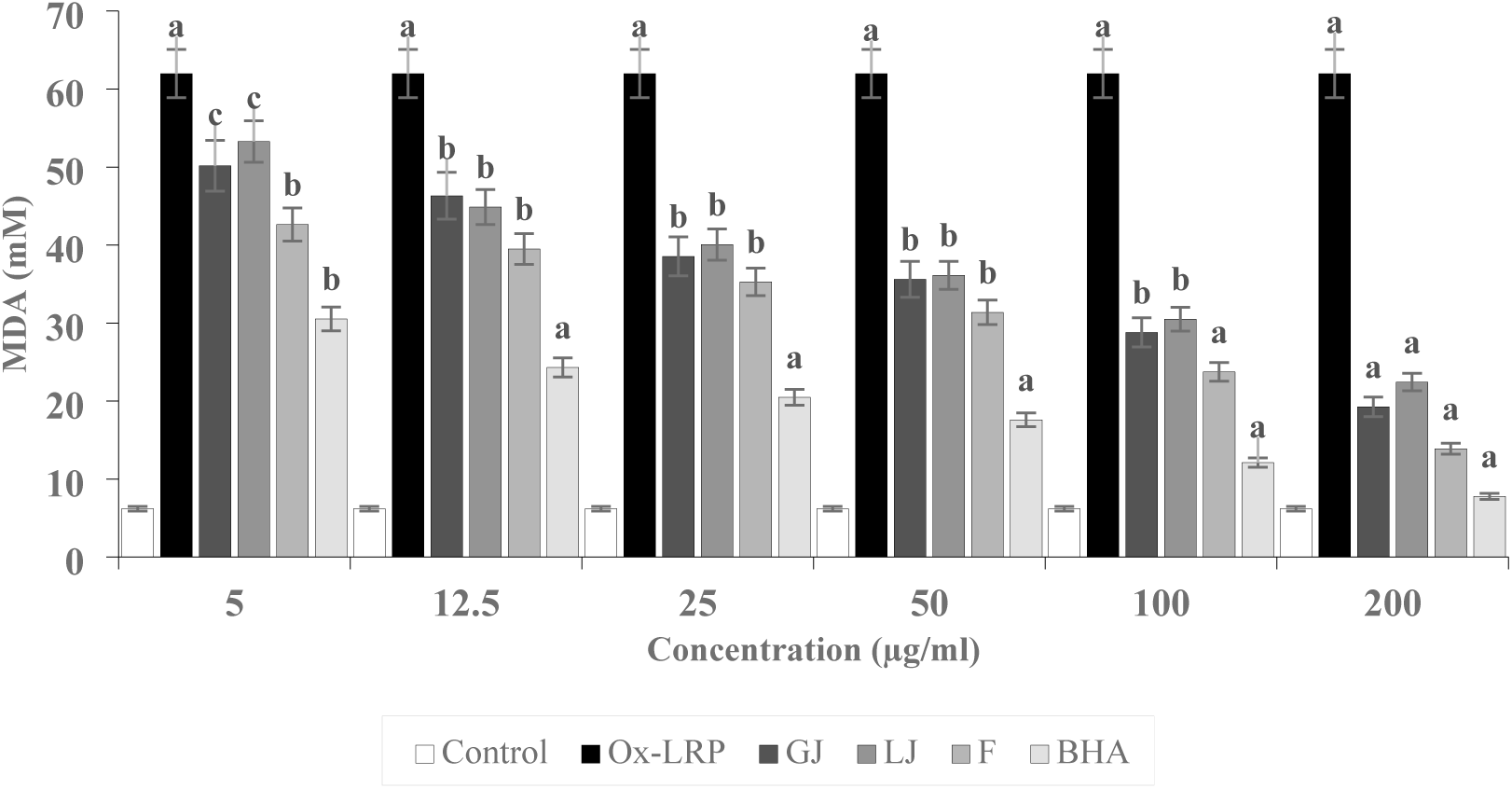
Antioxidant Effects of *Zingiber officinale*, *Citrus limon*, and Their Formulation on Lipid Peroxidation in Triton-Induced Lipid-Rich Plasma Assessed by TBARS Assay The ox-LRP group was compared with the Control group, and the GJ, LJ, F, and BHA groups were compared with the ox-LRP group. Values with different superscripts (a, b, or c) significantly differ at p < 0.05 (c), p < 0.01 (b), p < 0.001 (a)

The treatment with GJ demonstrated a significant, dose-dependent reduction in MDA levels, indicating the potent antioxidant properties of *Zingiber officinale*. This effect becomes particularly pronounced at higher concentrations (100–200 µg/mL), suggesting that the bioactive compounds present in ginger, such as gingerols and shogaols, play a crucial role in mitigating lipid peroxidation. Similarly, LJ displayed a comparable pattern of antioxidant activity, with a notable decrease in MDA levels, especially at concentrations of 50 µg/mL and above. The effectiveness of LJ can be attributed to its high content of polyphenolic compounds and flavonoids, which are known for their ability to scavenge free radicals and stabilize cellular lipids (figure 5).

The formulation (F), combining GJ and LJ, exhibited an enhanced antioxidant effect compared to the individual juices at corresponding concentrations. This observation suggests a potential synergistic interaction between the phytochemical compounds of ginger and lemon, where their combined antioxidant mechanisms surpass the sum of their individual effects. The reduction in MDA levels in response to the formulation (F) at the highest concentrations approached that observed with BHA, a synthetic antioxidant known for its robust inhibitory effect on lipid peroxidation. This result underscores the potential of the combined natural extracts as an effective alternative to synthetic antioxidants, providing comparable protection against oxidative stress (figure 5).

The BHA group, serving as a reference for maximal antioxidant protection, maintained near-baseline MDA levels across all concentrations, confirming the sensitivity and reliability of the TBARS assay in detecting lipid peroxidation inhibition. Statistical analysis, indicated by distinct superscript letters, validates the significant differences observed between the experimental groups, confirming the reproducibility and relevance of the results. The overall findings strongly support the hypothesis that the synergistic formulation of *Zingiber officinale* and *Citrus limon* offers superior antioxidant protection compared to individual treatments, reinforcing the potential of this combination as a natural therapeutic strategy to combat oxidative stress and related metabolic disturbances (figure 5).

### 3.5. Hypolipidemic effect of *Zingiber officinale* and *Citrus limon* juices on mice

The results of hypolipidemic activity are represented in **Figure 6**. Triton injection significantly increased triglycerides, low-density lipoprotein (LDLc), and total cholesterol while considerably lowering high-density lipoprotein (HDLc) when compared to average values (p < 0.05) **(figure 6-A to Figure 6-D**).

**Figure 6.**
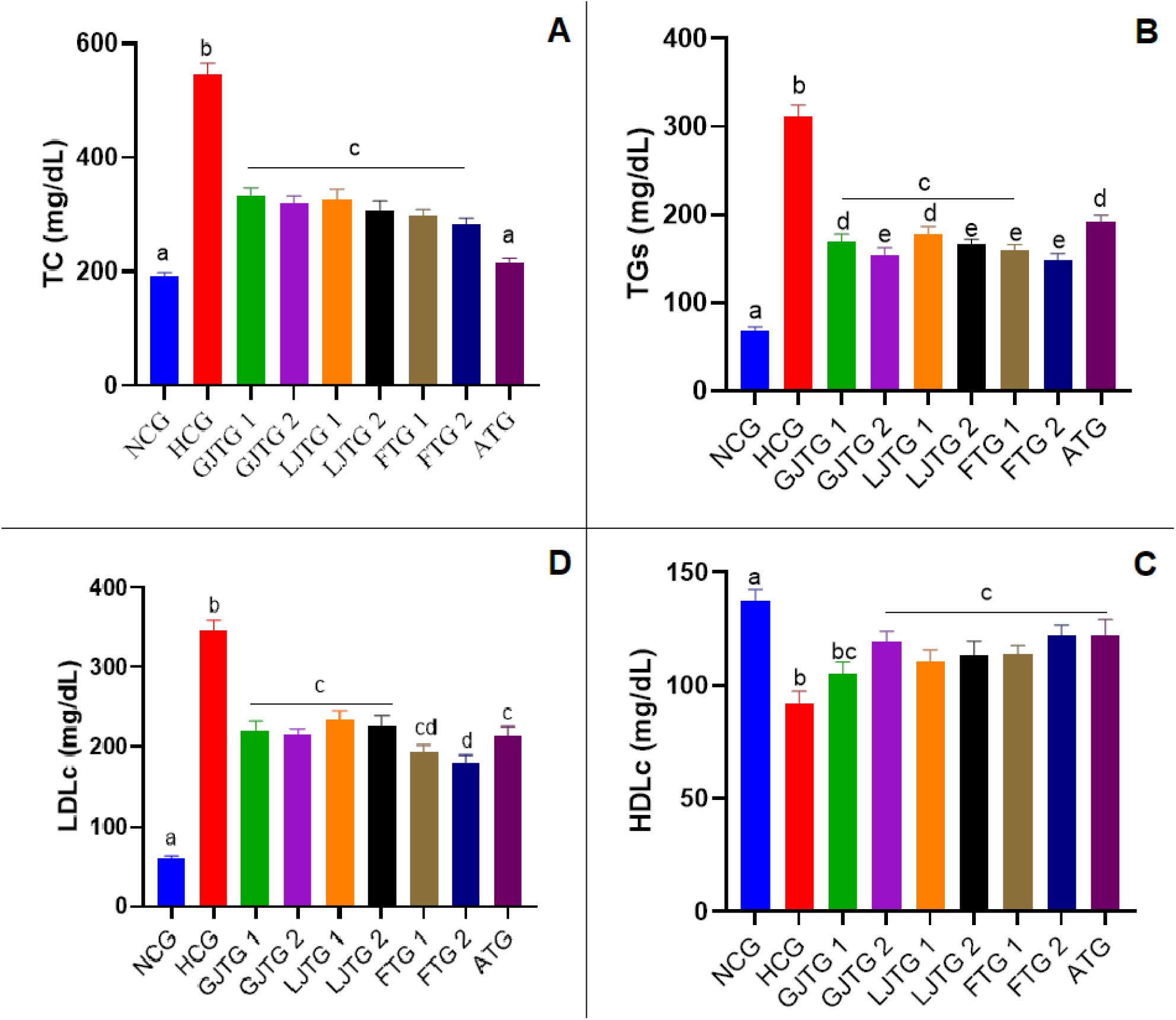
Effect of different *Zingiber officinale* and *Citrus limon* juices on plasmatic total cholesterol (**A**), triglycerides (**B**), HDLc (**C**), and LDLc (**D**) in acute dyslipidemic mice. NCG: Normal control group; HCG: Hyperlipidemic group; GJTG 1: Ginger juice extract treated group at 250 mg/Kg; GJTG 2: Ginger juice treated group at 500 mg/Kg; LJTG 1: Lemon juice treated group at 250 mg/Kg; LJTG 2: Lemon juice treated group at 500 mg/Kg; ATG: Atorvastatin treated group at 10 mg/Kg; FTG 1: Formulation treated group at 250 mg/Kg; FTG 2: Formulation treated group at 500 mg/Kg; The HCG group was compared with the NCG group, and the GJTG 1, GJTG 2, LJTG 1, LJTG 2, and ATG groups are compared with the HCG group. Values with different superscripts (a, b, c, d, or e) significantly differ at p < 0.05.

However, all treatments induced significant lipid-lowering effects compared to the hyperlipidemic group (p < 0.05). Indeed, administering juices at all concentrations and their combinations reduced elevated lipids with a comparable impact to Atorvastatin (10 mg/kg). Interestingly, at 500 mg/kg, the *Zingiber officinale* and *Citrus limon* juice formulation revealed higher TGs and LDL lowering effect (p < 0.05), suggesting synergetic effects.

#### AIP, LDL/HDL, TC/HDL, and TG/HDL ratios calculation

The atherogenic index of plasma (AIP) provides a valuable measure of the balance between atherogenic lipoproteins (such as LDL) and anti-atherogenic HDL particles. This index strongly predicts cardiovascular risk, with higher values correlating with increased lipid deposition in arterial walls and a higher propensity for atherogenesis. In the control group (NCG), the AIP is low, approximately 0.43, indicating a healthy lipid profile with balanced cholesterol fractions and minimal cardiovascular risk. This reflects a normal physiological state where HDL-C efficiently counters the potential atherogenic effects of total cholesterol. Conversely, in the hyperlipidemic control group (HCG), the AIP increases dramatically to 5.00. This sharp rise signifies severe dyslipidemia, characterized by a depletion of HDL-C and a substantial elevation of total cholesterol, indicative of an unregulated and highly atherogenic lipid environment caused by Triton WR-1339 administration (table 4).

**Table 4.** Effects of *Zingiber officinale* and *Citrus limon* juices on lipid ratios and Atherogenic Index in hyperlipidemic mouse models.

| Group | AIP | LDL/HDL | TC/HDL | TG/HDL |
| --- | --- | --- | --- | --- |
| NCG | 0.428 ± 0.02 | 0.642 ± 0.01 | 1.428 ± 0.05 | 0.714 ± 0.03 |
| HCG | 5.018 ± 0.10<sup>a</sup> | 3.202 ± 0.15<sup>a</sup> | 6.011 ± 0.20<sup>a</sup> | 3.005 ± 0.10<sup>a</sup> |
| GJTG1 | 3.090 ± 0.08<sup>b</sup> | 2.025 ± 0.12<sup>c</sup> | 4.090 ± 0.15<sup>b</sup> | 1.818 ± 0.08<sup>a</sup> |
| GJTG2 | 2.333 ± 0.07<sup>a</sup> | 1.666 ± 0.10<sup>a</sup> | 3.333 ± 0.12<sup>a</sup> | 1.529 ± 0.07<sup>a</sup> |
| LJTG1 | 2.565 ± 0.06<sup>a</sup> | 1.826 ± 0.09<sup>b</sup> | 3.565 ± 0.10<sup>b</sup> | 1.652 ± 0.06<sup>a</sup> |
| LJTG2 | 2.253 ± 0.05<sup>a</sup> | 1.666 ± 0.08<sup>a</sup> | 3.256 ± 0.09<sup>a</sup> | 1.538 ± 0.05<sup>a</sup> |
| FTG1 | 2.041 ± 0.04<sup>a</sup> | 1.524 ± 0.07<sup>a</sup> | 3.041 ± 0.08<sup>a</sup> | 1.366 ± 0.04<sup>a</sup> |
| **FTG2** | 1.846 ± 0.03<sup>a</sup> | 1.384 ± 0.06<sup>a</sup> | 2.846 ± 0.07<sup>a</sup> | 1.230 ± 0.03<sup>a</sup> |
| **ATG** | 1.592 ± 0.02<sup>a</sup> | 1.259 ± 0.05<sup>a</sup> | 2.592 ± 0.06<sup>a</sup> | 1.111 ± 0.02<sup>a</sup> |

In the treatment groups, a progressive reduction in AIP values is observed, demonstrating the efficacy of the interventions in mitigating the atherogenic burden. For instance, in the GJTG1 group, the AIP is reduced to 3.09, while further decreases are seen in groups such as FTG2 (2.33) and ATG (1.59). The almond treatment (ATG) exhibits the most significant improvement, reflecting its potential to elevate HDL-C levels or lower total cholesterol concentrations, thereby restoring lipid homeostasis. These findings underscore the therapeutic potential of the tested treatments in reducing cardiovascular risk by shifting the lipid balance toward a less atherogenic profile (table 4).

The LDL/HDL ratio is another critical marker, reflecting the proportion of low-density lipoprotein cholesterol (LDL-C) to high-density lipoprotein cholesterol (HDL-C). Higher values indicate an increased cardiovascular risk due to a more significant presence of atherogenic LDL particles relative to protective HDL particles. The LDL/HDL ratio in the NCG group is low (approximately 0.64), consistent with a low-risk lipid profile. In contrast, the HCG group exhibits a significantly elevated ratio of 3.2, confirming severe dyslipidemia and heightened cardiovascular risk. The treatment groups show notable reductions in this ratio, with ATG demonstrating the most pronounced effect, reducing the ratio to approximately 1.26. This suggests that the treatments effectively enhance the lipid profile by lowering LDL-C or increasing HDL-C, thus decreasing the relative burden of atherogenic particles. The TC/HDL ratio is widely regarded as a robust predictor of cardiovascular risk, offering insight into the overall balance of cholesterol fractions. A ratio below 3.5 is generally associated with a low risk of cardiovascular disease, while higher ratios indicate an increased risk. The NCG group’s ratio is 1.43, reflecting excellent cholesterol management. However, in the HCG group, this ratio rises drastically to 6.00, confirming the atherogenic impact of Triton WR-1339. Treatment groups show gradual improvement, with reductions in the ratio correlating with the efficacy of the interventions. The ATG group achieves the lowest ratio among the treated groups (approximately 2.59), suggesting a robust protective effect against lipid imbalance (table 4).

The TG/HDL ratio, a measure of triglyceride burden relative to HDL-C, is particularly relevant for assessing metabolic health and insulin resistance. A high TG/HDL ratio is a hallmark of dyslipidemia and a precursor to cardiovascular complications. In the NCG group, this ratio is approximately 0.71, indicative of a healthy lipid profile. The HCG group, however, shows a significant increase to 3.00, highlighting the pro-dyslipidemic effects of Triton WR-1339. With treatment, this ratio improves substantially, with ATG again demonstrating the most remarkable improvement, reducing the ratio to 1.11. This finding suggests that the therapy alleviates hypercholesterolemia and addresses hypertriglyceridemia, further supporting its therapeutic potential (table 3).

### 3.6. Toxicity study of *Zingiber officinale* and *Citrus limon* juices

#### 3.6.1. Acute toxicity study

The acute oral toxicity test demonstrated that the treated mice displayed normal behavior throughout the observation period of 14 days, even at the highest administered dose of 10 g/kg body weight. No signs of toxicity, such as changes in physical appearance, behavior, or other adverse effects, were observed, and no weight loss. This lack of harmful effects, even at elevated doses, indicates that GJ, LJ, and formulation F are safe and non-toxic under the experimental conditions of this study. These findings support their potential for further application without immediate safety concerns.

#### 3.6.2. Subacute toxicity study

##### a) Body weight

Figure 6 shows the graph representing the findings of a subacute toxicity study conducted over 28 days to evaluate the effects of *Zingiber officinale* (ginger juice), *Citrus limon* (lemon juice), and their combined formulation on rat body weights. The experiment included a Control and ten treated groups (G1 to G9). The Control group likely received no treatment or a placebo, while the treated groups received varying doses or formulations of the juices. Body weight was regularly measured on the 1^st^, 7^th^, 14^th^, 21^st^, and 28^th^ days to monitor growth patterns and detect potential toxic effects. The rats were kept under standard laboratory conditions, including controlled temperature, proper nutrition, and access to water, ensuring minimal environmental interference with the results.

From the graph, the body weights of all groups, including the Control and treated groups, show a general upward trend throughout the 28-day study period.

On the 1^st^ day, the body weights were nearly identical across all groups, indicating proper randomization and baseline uniformity before treatment initiation.

By the 7^th^ day, a slight increase in body weight is observed in all groups, including the control group and G1 to G9. However, the control group consistently demonstrated a higher body weight increase than the treated groups (figure 7).

**Figure 7.**
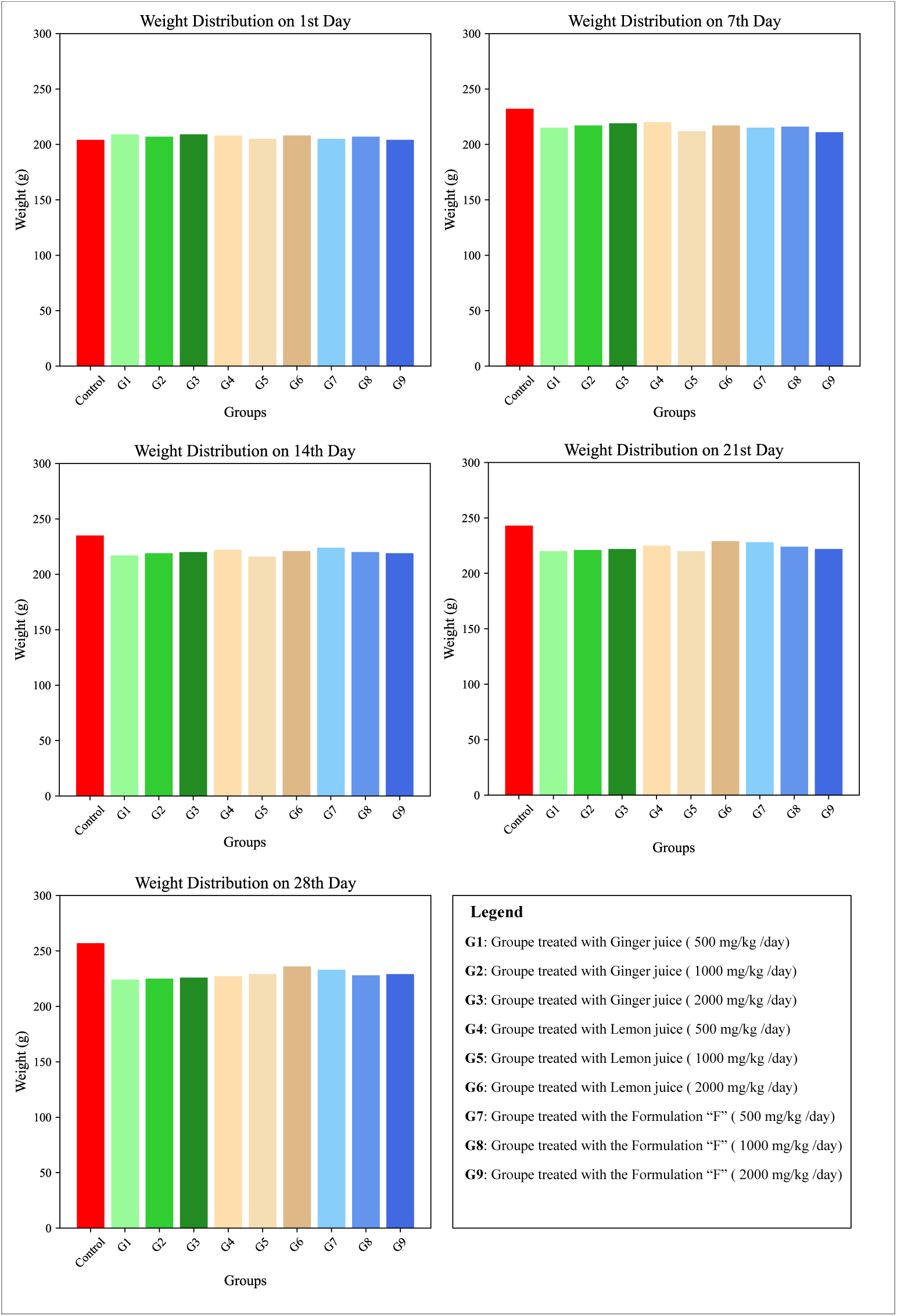
Body Weight variations in rats treated with ginger juice, lemon juice, and their formulation during a 28-day subacute toxicity study (Data are expressed as mean values ± standard deviation

By the 14^th^, 21^st^, and 28^th^ days, the body weights of the treated groups (G1 to G9) continue to increase but remain slightly lower than the Control group, with the difference being most noticeable on the 21st and 28th days. Despite this, the overlapping error bars suggest that the observed differences are not statistically significant (figure 7).

The steady increase in body weight across all groups, without any signs of stagnation or decline, indicates that the treatments did not induce severe systemic toxicity.

The body weight trends observed over the 28 days suggest that administering *Zingiber officinale*, *Citrus limon*, and their combined formulation did not result in acute or severe toxicity. Body weight is a critical indicator of systemic health in toxicity studies, and a consistent increase in weight across all groups supports the safety of these treatments.

The slightly lower weight gain in treated groups than in the control could indicate a minor modulatory effect on appetite, metabolism, or nutrient absorption. This is consistent with the known bioactive compounds in ginger and lemon, such as gingerols, citric acid, and flavonoids, which can influence digestion and metabolic processes. However, the absence of weight loss or growth suppression highlights that the dosages were within a safe range and did not disrupt overall growth and health (figure 7).

##### b) Relative organ weights

Table 4 provides data on the absolute and relative organ weights of rats subjected to oral administration of ginger juice (GJ), lemon juice (LJ), and their combined formulation (F) across ten experimental groups.

The absolute liver weight in the control group (G1) was 7.79 ± 0.51 g, with comparable values across all treatment groups. Notably, the highest absolute liver weight was observed in G3 (8.05 ± 0.49 g), while the lowest was recorded in G6 (7.85 ± 0.63 g). These results indicate a general consistency in liver weights across the treatments, with no significant deviations from the control group.

Relative liver weights exhibited a similar trend, with the control group showing a value of 3.31 ± 0.19 g. Minimal variations were observed among the treated groups, with G2 and G9 showing slightly higher values (3.40 ± 0.20 g and 3.51 ± 0.26 g, respectively), while G5 displayed one of the lower values (3.38 ± 0.18 g). These variations remain within a narrow range, suggesting that the treatments did not significantly influence liver weight relative to body weight (table 5).

**Table 5.** Absolute and relative organ weights of rats administered orally with the GJ, LJ, and their formulation F.

|  | Parameters | Control | G1 | G2 | G3 | G4 | G5 | G6 | G7 | G8 | G9 |
| --- | --- | --- | --- | --- | --- | --- | --- | --- | --- | --- | --- |
| Liver (g) | Absolute weight (g) | 7.79 ± 0.51 | 7.88 ± 0.40 | 8.05 ± 0.49 | 7.96 ± 0.63 | 7.99 ± 0.63 | 7.85 ± 0.63 | 7.91 ± 0.63 | 8.11 ± 0.63 | 7.76 ± 0.63 | 8.02 ± 0.50 |
|  | Relative weight (g) | 3.31 ± 0.19 | 3.44 ± 0.22 | 3.40 ± 0.20 | 3.49 ± 0.28 | 3.46 ± 0.21 | 3.38 ± 0.18 | 3.45 ± 0.27 | 3.44 ± 0.28 | 3.40 ± 0.25 | 3.51 ± 0.26 |
| Kidney (g) | Absolute weight (g) | 0.70 ± 0.01 | 0.68 ± 0.02 | 0.65 ± 0.01 | 0.66 ± 0.02 | 0.63 ± 0.02 | 0.61 ± 0.01 | 0.64 ± 0.02 | 0.70 ± 0.02 | 0.66 ± 0.01 | 0.69 ± 0.02 |
|  | Relative weight (g) | 0.25 ± 0.01 | 0.27 ± 0.01 | 0.25 ± 0.01 | 0.26 ± 0.01 | 0.27 ± 0.01 | 0.25 ± 0.01 | 0.26 ± 0.01 | 0.28 ± 0.01 | 0.24 ± 0.01 | 0.29 ± 0.01 |

Data are presented as Mean ± SEM; **G1**: control group treated with distilled water; **G2**: group treated with 500 mg/kg/day of GJ; **G3**: group treated with 1000 mg/kg/day of GJ; **G4**: group treated with 2000 mg/kg/day of GJ; **G5**: group treated with 500 mg/kg/day of LJ; **G6**: group treated with 1000 mg/kg/day of LJ; **G7**: group treated with 2000 mg/kg/day of LJ; **G8**: group treated with 500 mg/kg/day of F; **G9**: group treated with 1000 mg/kg/day of F; **G10**: group treated with 2000 mg/kg/day of F.

The absolute values in the control group for kidney weights were measured at 0.70 ± 0.01 g. Across the treatment groups, absolute kidney weights ranged from 0.61 ± 0.01 g in G5 to 0.70 ± 0.02 g in G7. While there were slight fluctuations, particularly in G6 (0.64 ± 0.02 g) and G8 (0.66 ± 0.01 g), these differences were minimal and unlikely to indicate significant biological effects. Relative kidney weights were consistent across all groups, with the control group at 0.25 ± 0.01 g and values ranging from 0.24 ± 0.01 g to 0.29 ± 0.01 g among the treated groups. The highest relative kidney weight was recorded in G9 (0.29 ± 0.01 g), corresponding to the highest dose of the formulation, but this increase remained modest (table 5).

In general, the comparison of organ weights across the experimental groups highlights the negligible impact of the treatments on both absolute and relative organ weights. The administration of GJ, LJ, and their formulation F, even at the highest doses, did not result in substantial deviations from the control group, indicating that these treatments are well-tolerated. The stability of both liver and kidney weights across groups supports the conclusion that the tested substances do not induce organ toxicity under the conditions of this study.

##### c) Hematological analysis

Table 5, in turn, illustrates the effects of subacute oral administration of ginger juice (GJ), lemon juice (LJ), and their combined formulation (F) on the hematological parameters of rats. The data, expressed as mean ± SEM, compares results across ten groups, from groups 1 (G1) to 9 (G9).

To begin with, white blood cell (WBC) counts showed minor fluctuations among groups, with values ranging from 4.76 ± 0.14 × 10⁹/L in G9 to 5.13 ± 0.13 × 10⁹/L in G7, compared to 4.95 ± 0.13 × 10⁹/L in the control group. These findings suggest minimal impact of the treatments on immune cell counts. Similarly, red blood cell (RBC) levels were consistent across treatments, with the highest value recorded in G5 (7.20 ± 0.37 × 10¹²/L) and the lowest in G4 (6.85 ± 0.35 × 10¹²/L). The control group reported an RBC count of 6.99 ± 0.31 × 10¹²/L, and treated groups, including those receiving the combined formulation (F), demonstrated stable RBC values (table 6).

**Table 6.** Impact of subacute oral administration of GJ, LJ, and their formulation F on the hematological profile of rats.

|  | WBC (10<sup>9</sup>/L) | RBC (10<sup>12</sup>/L) | HGB (g/dL) | HCT (%) | MCV (fL) | MCH (pg) | MCHC (g/dL) | PLT (10<sup>9</sup>/L) | MPV (fL) |
| --- | --- | --- | --- | --- | --- | --- | --- | --- | --- |
| **Control** | 4.95 ± 0.13 | 6.99 ± 0.31 | 12.96 ± 0.49 | 38.06 ± 0.42 | 54.15 ± 0.74 | 17.96 ± 0.60 | 35.09 ± 0.57 | 602.66 ± 17.83 | 6.49 ± 0.22 |
| **G1** | 5.12 ± 0.16 | 7.17 ± 0.34 | 13.09 ± 0.46 | 38.11 ± 0.46 | 54.18 ± 0.72 | 18.03 ± 0.58 | 34.91 ± 0.59 | 598.17 ± 21.58 | 6.52 ± 0.20 |
| **G2** | 4.89 ± 0.19 | 7.11 ± 0.32 | 13.11 ± 0.51 | 37.96 ± 0.35 | 53.94 ± 0.70 | 18.11 ± 0.49 | 35.07 ± 0.51 | 611.09 ± 20.99 | 6.53 ± 0.22 |
| **G3** | 4.91 ± 0.17 | 6.95 ± 0.38 | 13.12 ± 0.54 | 38.13 ± 0.33 | 54.07 ± 0.73 | 18.14 ± 0.61 | 35.15 ± 0.56 | 609.28 ± 24.15 | 6.44 ± 0.26 |
| **G4** | 5.03 ± 0.13 | 6.85 ± 0.35 | 12.93 ± 0.46 | 37.98 ± 0.29 | 53.93 ± 0.68 | 18.12 ± 0.53 | 35.12 ± 0.53 | 599.53 ± 20.47 | 6.45 ± 0.31 |
| **G5** | 4.97 ± 0.15 | 7.20 ± 0.37 | 13.06 ± 0.48 | 37.90 ± 0.50 | 54.12 ± 0.74 | 17.95 ± 0.48 | 35.26 ± 0.48 | 608.87 ± 22.70 | 6.39 ± 0.31 |
| **G6** | 4.90 ± 0.19 | 7.15 ± 0.42 | 12.98 ± 0.59 | 37.92 ± 0.34 | 53.91 ± 0.71 | 18.13 ± 0.64 | 34.98 ± 0.52 | 596.38 ± 19.87 | 6.48 ± 0.25 |
| **G7** | 5.13 ± 0.13 | 7.14 ± 0.36 | 13.15 ± 0.61 | 38.25 ± 0.48 | 54.03 ± 0.68 | 17.90 ± 0.57 | 35.18 ± 0.51 | 598.05 ± 25.04 | 6.43 ± 0.29 |
| **G8** | 4.99 ± 0.12 | 6.92 ± 0.43 | 13.18 ± 0.50 | 37.97 ± 0.47 | 54.01 ± 0.70 | 18.05 ± 0.59 | 35.12 ± 0.44 | 604.54 ± 16.08 | 6.56 ± 0.27 |
| **G9** | 4.76 ± 0.14 | 6.89 ± 0.40 | 12.99 ± 0.45 | 38.17 ± 0.31 | 53.98 ± 0.71 | 17.98 ± 0.62 | 35.17 ± 0.50 | 602.21 ± 23.65 | 6.51 ± 0.30 |

Values are expressed as Mean ± SEM; **G1**: group treated with 500 mg/kg/day of GJ; **G2**: group treated with 1000 mg/kg/day of GJ; **G3**: group treated with 2000 mg/kg/day of GJ; **G4**: group treated with 500 mg/kg/day of LJ; **G5**: group treated with 1000 mg/kg/day of LJ; **G6**: group treated with 2000 mg/kg/day of LJ; **G7**: group treated with 500 mg/kg/day of F; **G8**: group treated with 1000 mg/kg/day of F; **G9**: group treated with 2000 mg/kg/day of GJ; WBC: white blood cell count; RBC: red blood cell count; HGB: hemoglobin concentration; HCT: hematocrit; MCV: mean corpuscular volume; MCH: mean corpuscular hemoglobin; MCHC: mean corpuscular hemoglobin concentration; PLT: platelet levels; MPV: mean platelet volume.

**Table 6.** Chemical composition of *Zingiber officinale* and *Citrus limon* juices.

| Juice | Compound |
| --- | --- |
| *Z. officinale* | 4-gingerol |
|  | 6-gingediol |
|  | 6-gingerol |
| Juice | Compound |
| *C. limon* | Eriodictyol |
|  | Rutin |
|  | Hesperidin |
|  | Isorhamnetin |

Furthermore, hemoglobin (HGB) concentrations showed slight variations across the groups, with the highest level in G8 (13.18 ± 0.50 g/dL) and the lowest in G4 (12.93 ± 0.46 g/dL). The control group (12.96 ± 0.49 g/dL) remained within a similar range to the treated groups, reflecting no significant alterations. In addition, hematocrit (HCT) values followed a similar trend, with the control group at 38.06 ± 0.42% and the other groups exhibiting values between 37.96 ± 0.35% and 38.25 ± 0.48%, highlighting the stability of this parameter across all treatments (table 6).

Moreover, erythrocyte indices, including mean corpuscular volume (MCV), mean corpuscular hemoglobin (MCH), and mean corpuscular hemoglobin concentration (MCHC), were broadly consistent among groups. The MCV values ranged from 53.91 ± 0.68 fL in G6 to 54.18 ± 0.72 fL in G5, with the control group at 54.15 ± 0.74 fL. Likewise, the MCH values showed minimal variation, ranging from 17.95 ± 0.48 pg in G5 to 18.11 ± 0.49 pg in G2. Similarly, the MCHC values ranged from 34.91 ± 0.59 g/dL in G1 to 35.26 ± 0.48 g/dL in G5, comparable to the control group’s 35.09 ± 0.57 g/dL value (table 6).

Finally, platelet counts (PLT) were stable across all groups, with the highest count observed in G5 (608.87 ± 22.70 × 10⁹/L) and the lowest in G6 (596.38 ± 19.87 × 10⁹/L). The control group reported 602.66 ± 17.83 × 10⁹/L, and groups treated with the formulation F, such as G7 (604.54 ± 16.08 × 10⁹/L) and G10 (602.21 ± 23.65 × 10⁹/L), displayed no significant deviations. Mean platelet volume (MPV) values were consistent across all groups, ranging from 6.39 ± 0.31 fL in G5 to 6.56 ± 0.27 fL in G8, with the control group showing a value of 6.49 ± 0.22 fL (table 6).

In conclusion, the hematological parameters across all groups remained within a narrow range of variation, indicating that oral administration of GJ, LJ, and their formulation F had no significant adverse effects on blood cell counts, hemoglobin levels, or platelet profiles. These findings suggest a stable hematological response to the treatments.

### 3.7. Molecular Docking Study

Molecular priming is a highly effective mechanism, providing insights into potential interactions between a molecule and its targeted protein receptor, inferred from the energy discharge during molecular engagements (25–27). This technique is a foundational computational tool frequently used to delineate the critical molecular interactions of pharmacologically active entities.

Figure 8 shows the structure of one of the main enzymes implicated in lipid metabolism, HMG-CoA reductase.

**Figure 8.**
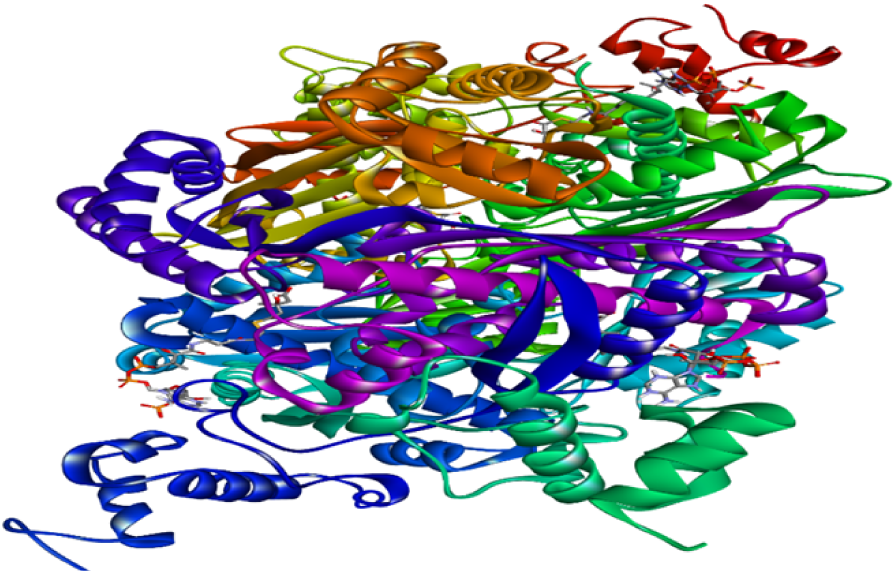
3D structure of the HMG-CoA reductase.

Additionally, it can be perceived as a preliminary step in the developmental protocol for novel therapeutic agents. The present study conducted priming analyses on four molecules sourced from *C. limon* juice and three from *Z. officinale* juice (table 4) in conjunction with the enzyme.

The binding energy varied according to the molecule (table 4), the lowest being that of the bond between “HMG-CoA reductase” (figure 7) and “6-gingerol” (table 6) and the highest being that of the bond between “HMG-CoA reductase” and “hesperidin.” It even has higher energy than the “HMG-CoA reductase” inhibitor, which means that it has a very high probability of inhibiting “HMG-CoA reductase”, which means that in the case of hesperidin, there is a very high chance of inhibiting “HMG-CoA reductase” and thus decreasing the synthesis of endogenous cholesterol, the high affinity of hesperidin is due to intramolecular solid interactions with the active site of the enzyme because of hydrogen bonds between the oxygen of the carbonyl group with the residue LYS C:633, HIS C:536 the hydrogen bonds between the oxygen of the hydroxyl group with the residues Leu B: 634, ILE B:699, SER C;637, GLU B:700, SER B:705, GLN B:648, HIS C:635, LYS C633 the high affinity is also due to the presence of other types of interactions such as alkyl or pi-alkyl type interactions or pi-cation type interactions (figure 9).

**Figure 9.**
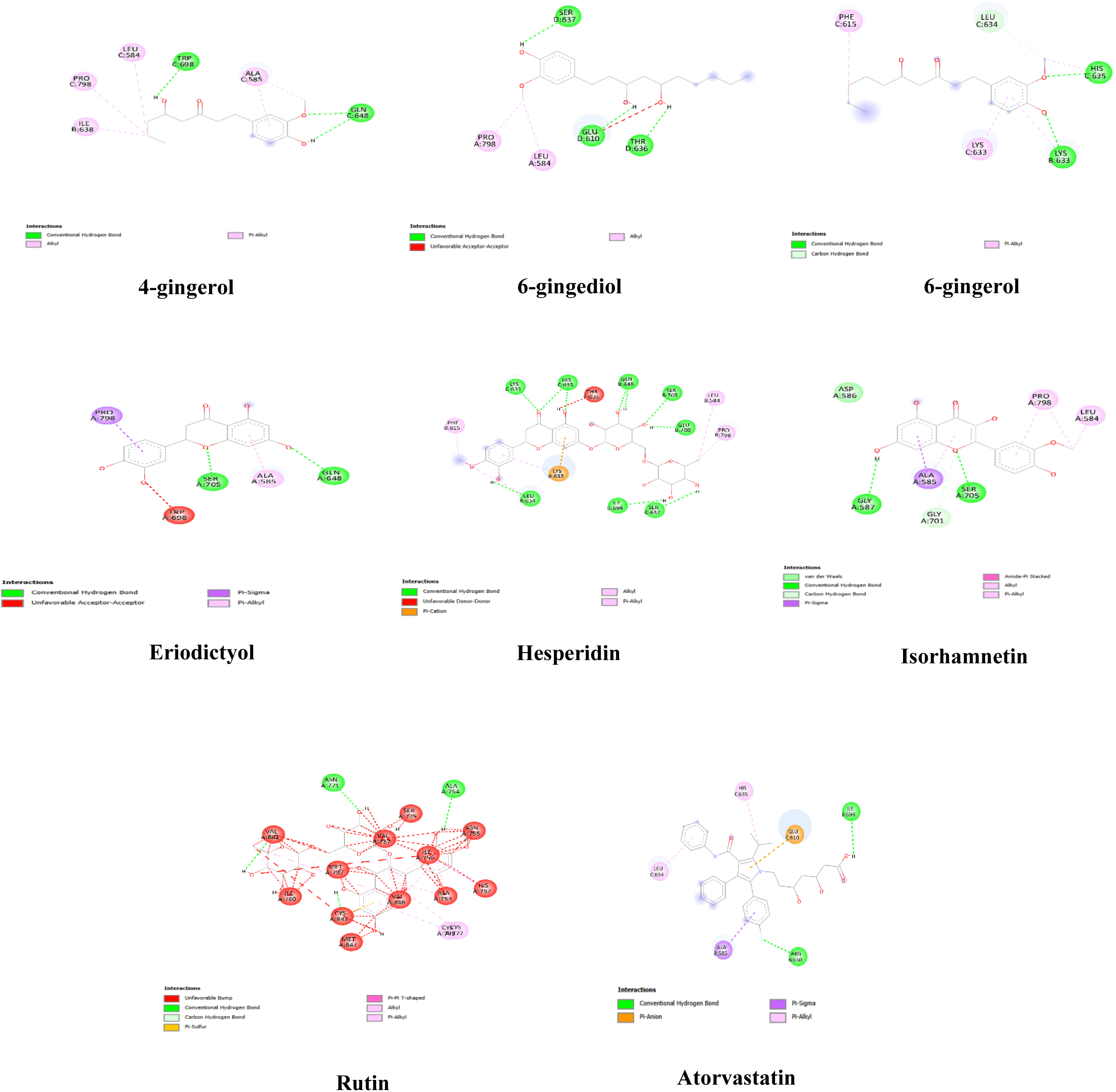
Possible interaction sites of *Zingiber officinale*, *Citrus limon,* and the atorvastatin molecules.

Table 7 provides an insightful comparison of the binding energies of several natural compounds derived from ginger and lemon and a reference drug, Atorvastatin, with the HMG-CoA reductase enzyme. As a critical regulator of cholesterol biosynthesis, HMG-CoA reductase is a well-established therapeutic target in lipid metabolism. The data presented here allow us to evaluate the potential of these natural compounds as inhibitors of this enzyme based on their binding affinities. Lower (more negative) binding energy values indicate stronger interactions and a higher likelihood of effective inhibition.

**Table 7.** The binding energy of the molecules of *Zingiber officinale* and *Citrus limon* molecules with HMG-CoA reductase.

| Molecule | Binding Energy (Kcal/mol) |
| --- | --- |
| Rutin | -9 |
| Hesperidin | -10.4 |
| Eriodictyol | -8.5 |
| Isorhamnetin | -8 |
| 6-gingerol | -5.5 |
| 6-gingediol | -6.1 |
| 4-gingerol | -6.1 |
| Atorvastatin | -8.1 |

Among the compounds analyzed, hesperidin demonstrates the most potent interaction with the enzyme, with a binding energy of -10.4 Kcal/mol, surpassing the value for Atorvastatin (-8.1 Kcal/mol), a widely used cholesterol-lowering drug. This remarkable result positions hesperidin as a promising natural alternative with significant potential to modulate lipid levels. Similarly, rutin (-9 Kcal/mol) and eriodictyol (-8.5 Kcal/mol) show strong binding affinities, suggesting they might also be valuable candidates for further exploration. Other compounds, such as isorhamnetin (-8 Kcal/mol), display slightly lower affinity but still hold potential therapeutic interest. On the other hand, molecules derived from ginger, including 6-gingerol (-5.5 Kcal/mol), 6-gingediol (-6.1 Kcal/mol), and 4-gingerol (-6.1 Kcal/mol), exhibit more moderate binding energies, indicating a less pronounced interaction with the enzyme.

These findings are particularly noteworthy when compared to Atorvastatin, a benchmark in cholesterol management. Specific natural molecules, especially Hesperidin and Rutin, exhibit stronger binding affinities, highlighting their potential to serve as effective inhibitors. This could lead to the development of safer and possibly more accessible therapeutic options. Natural compounds often bring additional benefits, such as reduced side effects and improved tolerability, making them appealing for therapeutic use. However, it is essential to note that these preliminary results should be followed by in-depth investigations to confirm their biological activity and therapeutic potential.

The biological implications of these results are profound. These natural molecules could help regulate cholesterol levels and support cardiovascular health by inhibiting HMG-CoA reductase. The lower affinities observed with ginger-derived compounds suggest they may have complementary benefits or target other pathways, making them useful in combination therapies.

In conclusion, the table highlights the significant potential of natural compounds from *Zingiber officinale* and *Citrus limon* as HMG-CoA reductase inhibitors. Hesperidin stands out as the most promising candidate, outperforming Atorvastatin in binding affinity, which emphasizes the need for further research. These findings underscore the growing interest in natural products as effective and safe alternatives in cholesterol management, providing a foundation for future studies to improve public health through natural-based therapies.

## Discussion

Whatever the causes, the number of hyperlipidemic subjects in the world today is increasing, as are the associated health costs. Hyperlipidemia can cause numerous complications, such as atherosclerosis, cardiovascular and cerebrovascular infarction, heart disease, and stroke. (Gong et al., 2020). Indeed, high lipid levels increase the risk of cardiovascular disease by negatively modifying the lipid profile, leading to the death of at least 2.8 million adults each year. (Alberti et al., 2005). Hence, pursuing novel lipid-lowering agents that offer enhanced safety and efficacy gains significance. The induction of acute hyperlipidemia in animal models using Triton WR-1339, a nonionic detergent (tertiary polymer octylphenol-oxyethylated formaldehyde), has gained widespread use as a method to assess the potential of both natural and synthetic anti-hyperlipidemic agents. (Schurr et al., 1972) and to investigate cholesterol and triacylglycerol metabolism (Abdou et al., 2018). The inhibition effect of lipoprotein lipase activity is thought to be one factor in the accumulation of plasma lipids caused by this detergent (Scharwey et al., 2013).

Furthermore, following 24 hours post-triton injection, an initial surge of 2 to 3 times the baseline lipid levels occurs, which diminishes over the next 24 hours. This triton-induced hyperlipidemia experimental model furnishes a relatively rapid and uncomplicated approach for evaluating potential anti-hyperlipidemic medications. It presents a swift and uncomplicated supplementary assessment tool during the initial screening phase (Friedman and Byers, 1953). Thus, it is used in numerous scientific investigations into hyperlipidemia research, using different animal models, such as rats (Bouhlali et al., 2020), mice (Touiss et al., 2017), hamsters (Ramchoun et al., 2009), and guinea pigs (Elbouny Hamza, 2023).

Since ancient times, medicinal and aromatic plants have been used to treat several disorders, including cardiovascular disease (Chaachouay et al., 2022). Recent scientific literature and clinical investigations have substantiated the effectiveness of numerous medicinal plants and herbal formulations in enhancing the balanced regulation of lipid metabolism and mitigating cardiovascular complications. (Ama Moor et al., 2017; Mishra, P. R., Panda, P. K., Apanna, K. C., & Panigrahi, 2011; Munasinghe et al., 2001). Indeed, using phytoconstituents and natural products to treat hyperlipidemia and cardiovascular disease could be beneficial due to their safety and the absence of side effects such as those reported for synthetic lipid-lowering agents (Dujovne, n.d.; Munasinghe et al., 2001). Based on this overview, we aim to evaluate the anti-hyperlipidemic activity of the Zingiber officinale and Citrus limon juice.

The main advantages of this present work are, first, the evaluation of the anti-hyperlipidemic effect of the juice of both plants separately, as well as that of their combination (50% of *Zingiber officinale* juice and 50% of *Citrus limon*). Second, the use of fresh juices from these plants, which are very easy to obtain and safe for administration, unlike organic solvent extracts, and thirdly, the validation of the findings of the *in vivo* study by an *in sillico* investigation using the “auto-docking-vina” software. The “LJ” extract exhibited a polyphenol and flavonoid content equivalent to 25.23 ± 1.54 mg GA/g of extract and 12.75 ± 3.08 mg QE/g of extract, respectively. This same extract was abundant in eriodictyol, rutin, hesperidin, and isorhamnetin. On the other hand, the “GJ” extract showed values corresponding to 18.48 ± 1.14 mg GA/g of extract and 7.26 ± 2.05 mg QE/g of extract for the polyphenol and flavonoid content, respectively. Chemical analysis revealed the presence of 4-gingerol, 6-gingediol, and 6-gingerol. Our findings closely align with those of a study conducted by [40]. It was reported that *Zingiber officinale*, *Citrus limon*, and their formulation contain many phenolic compounds and flavonoids.

The antioxidant studies we found confirm that the *Zingiber officinale* and *Citrus limon* juices and their formulation have an antioxidant potential that aligns with and expands upon previous studies investigating the antioxidant properties of *Zingiber officinale* and *Citrus limon*. The IC_50_ values obtained in the DPPH and ABTS assays for ginger and lemon juices are consistent with those reported in earlier research. For instance, a study by Pacula et al. reported IC_50_ values for ginger extracts ranging between 35 and 45 µg/mL in the DPPH assay (Pacuła et al., 2017), which closely mirrors the IC_50_ of 38.91 ± 1.47 µg/mL observed in this study. Similarly, lemon juice exhibited comparable values to those described by Kumar et al., where IC_50_ values for Citrus limon extracts ranged from 40 to 50 µg/mL depending on the solvent used for extraction (Kumar, 2019). These parallels suggest that the antioxidant activities of these extracts are robust across varying experimental conditions and methodologies.

Interestingly, the formulation of ginger and lemon juices did not consistently demonstrate enhanced antioxidant activity compared to the individual components. This finding contrasts with the work of Zhang et al., which showed a synergistic enhancement in antioxidant activity when combining ginger and citrus extracts (Zhang, 2020). However, it is worth noting that variations in the ratio of the combined extracts and the presence of other bioactive compounds, such as phenolics and flavonoids, may influence the overall antioxidant potential. Further studies investigating optimal blending ratios and the role of specific active compounds could provide a deeper understanding of these interactions. The TAC and FRAP assays in this study revealed relatively uniform antioxidant capacities among the samples, which aligns with findings by Antoniewicz et al., who highlighted the strong ferric-reducing power and total antioxidant capacities of *Zingiber officinale* and *Citrus limon* individually (Antoniewicz (Kałduńska) et al., 2021). However, the slightly reduced FRAP activity observed in the formulation may suggest the presence of antagonistic interactions between certain phytochemicals in the combined extracts. Such antagonistic effects have been previously described by Chen et al. in their work on polyphenol-rich plant combinations, emphasizing that not all mixtures yield synergistic effects (Chen, 2018).

Ascorbic acid served as a robust positive control in all assays, consistently demonstrating superior antioxidant activity compared to the natural extracts. This finding underscores the well-documented efficacy of ascorbic acid as a free radical scavenger and reducing agent, as reported by Kaur and Kapoor (Kaur and Kapoor, 2001). Nevertheless, the relatively close IC_50_ values of the natural extracts to ascorbic acid in the TAC and FRAP assays highlight their potential as alternative sources of antioxidants, particularly in functional food applications.

The variability in IC_50_ values between the DPPH and ABTS assays also deserves attention. This difference is consistent with prior studies that emphasize the distinct mechanisms of these assays. DPPH primarily measures the ability of antioxidants to donate hydrogen atoms to neutralize free radicals, while ABTS evaluates both hydrogen atom and electron transfer capabilities. This distinction may explain the marginal differences in antioxidant rankings observed across the assays.

Our findings highlight the strong antioxidant potential of *Zingiber officinale* (GJ), *Citrus limon* (LJ), and their combined formulation (F) in mitigating lipid peroxidation, as evidenced by significant reductions in MDA levels in the TBARS assay. The lipid-rich plasma model induced by Triton WR-1339 effectively simulated oxidative stress, allowing us to assess the protective effects of these natural extracts. GJ showed an apparent dose-dependent reduction in MDA levels, likely due to 4-gingerol, 6-gingerol, and 6-gingediol, known for their strong radical-scavenging properties and modulation of inflammatory pathways. Similarly, LJ exhibited significant antioxidant activity, particularly at higher concentrations, owing to its flavonoid content, including hesperidin, eriodyctiol, isorhamnetin, and rutin. These compounds have been shown to stabilize membranes, chelate metal ions, and inhibit lipid peroxidation. The combined formulation (F) demonstrated superior efficacy to the individual extracts, particularly at higher doses, indicating a synergistic effect. This likely results from the complementary actions of hydrophilic antioxidants, such as hesperidin, and lipophilic compounds, such as 6-gingediol, targeting both aqueous and lipid-phase oxidative reactions.

Our findings align with previous research highlighting the antioxidant properties of *Zingiber officinale* and *Citrus limon*. For instance, Bekkouch et al. (2022) demonstrated that ginger and lemon juice extracts, rich in compounds like 4-gingerol, 6-gingediol, hesperidin, and rutin, exhibit significant antioxidant potential and hepatoprotective effects against CCl₄-induced liver damage in rats (Bekkouch et al., 2022). Similarly, Köksal and Gülçin (2017) evaluated the antioxidant properties of ginger extracts, finding that ethanol extracts exhibited superior antioxidant capacity compared to water extracts, attributed to higher concentrations of phenolic acids such as ferulic acid and p-coumaric acid (Tohma et al., 2017).

Additionally, a study by Shende (2024) assessed the bioactive properties of aqueous and ethanolic extracts of *Citrus medica* and *Zingiber officinale*, revealing significant antioxidant, antibacterial, and anthelmintic effects (Shende, 2024), further supporting the therapeutic potential of these plants. These studies reinforce the efficacy of combining ginger and lemon extracts to achieve superior protection against oxidative damage, consistent with our observations. The fact that the formulation’s effect approached that of BHA, a potent synthetic antioxidant, reinforces its potential as a natural alternative for combating oxidative stress. Our findings align with existing studies on polyphenol-rich extracts but also highlight the unique strength of combining specific compounds, contributing to a broader protective effect and offering new insights into the potential for natural antioxidant formulations in managing oxidative damage.

On the other hand, our findings demonstrated that administering *Zingiber officinale* and *Citrus limon* juices attenuated the elevation of the primary plasma lipids. The lipid-lowering effects were comparable to Atorvastatin, one of the most prescribed medications for treating hyperlipidemia. Like *Zingiber officinale* and *Citrus limon*, several plants have been found to exhibit hypolipidemic properties (Khanna et al., 2002; Ramchoun et al., 2009). Two studies demonstrated that the polyphenol-rich extracts of basil and thyme reduced and retarded the synthesis of triglycerides and cholesterol (Khouya et al., 2021; Touiss et al., 2017). Similar conclusions were reported for phenolic bioactive phytochemicals in several herbal medicinal plants (Manzoni et al., 2019; Panahi et al., 2018; Rašković et al., 2019; Shen et al., 2019), in which it was found that they can prevent cholesterol absorption in enterocytes, decrease cholesterol synthesis, boost reverse cholesterol transport, and promote cholesterol excretion in the liver (Ibrahim et al., 2016).

Moreover, other lipid-lowering mechanisms were reported for polyphenols, including the activation of hepatic LDLc receptors responsible for the final clearance of cholesterol in the form of bile acids (Mbikay et al., 2014) and the alteration of cholesterol metabolism through the regulation of the activity and expression of critical enzymes such as HMG-CoA reductase (Tirawanchai et al., 2018), acyl CoA cholesterol acyl transferase (ACAT) (Cha et al., 2016), and lecithin cholesterol acyltransferase (LCAT) (Ama Moor et al., 2017). The richness of *Z. officinale* and *C. limon* juices in bioactive phenolic compounds can explain their remarkable lipid-lowering effect. Previous reports support the findings of our study (Kim and Ko, 2017; Murad et al., 2018; Oboh et al., 2015; Paul et al., 2013; Sharma et al., 1996). In our research, we have used a new extraction methodology, harnessing the potential of juice extracts to unlock valuable compounds without the need for conventional temperature-based or chemical solvent extraction techniques. Unlike traditional methods, often involving elevated temperatures and potentially harmful solvents, our approach signifies a paradigm shift towards a safer, more sustainable, and environmentally friendly extraction process. By exclusively relying on juice extracts, we preserve the target compounds’ inherent bioactive properties and eliminate the environmental impact associated with chemical solvents. This innovative methodology expands the scope of extraction techniques. It opens avenues for further exploration in natural product research, offering a unique and eco-friendly perspective on obtaining valuable bioactive components.

Our findings indicate that the elevated dosage (500 mg/kg) of the *Zingiber officinale* and *Citrus limon* juice blend exhibited a more pronounced reduction in lipid levels. This outcome is likely attributed to the synergistic effects stemming from the interactions between gingerols in *Zingiber officinale* juice and flavonols derived from *Citrus limon* juice. Numerous reports have demonstrated the potential of these compounds to alleviate hyperlipidemia. Specifically, isorhamnetin has been observed to possess the ability to facilitate macrophage injury induced by ox-LDL (oxidized low-density lipoprotein) (M et al., 2019) and attenuate atheromatous plaque formation by inhibiting macrophage apoptosis via PI3K/AKT activation and HO-1 induction in ox-LDL-treated THP-1-derived macrophages (Luo et al., 2015). Moreover, this bioactive flavonoid was also reported to induce a dose-dependent hypolipidemic activity in hyperlipidemic rats (NR et al., 2021). On the other hand, 6-gingerol exerted anti-hyperlipidemic effects in polaxamer P-407-induced hyperlipidemic rats (Shao et al., 2016). In addition, gingerol demonstrated an anti-hyperlipidemic effect in high-fat diet-induced hyperlipidemic rats by modulating the inflammatory pathways and the expression of enzymes essential to cholesterol metabolism, including lipoprotein lipase, LCAT, HMG-CoA reductase, and acetyl CoA carboxylase (Brahma Naidu et al., 2016).

HMG-CoA reductase regulates cholesterol production and limits the rate of the mevalonate pathway. (Ezeh and Ezeudemba, 2021a). The inhibition of the activity of this enzyme is one of the most effective treatments for reducing LDL levels, thereby preventing cardiovascular disease and atherosclerosis (Charan et al., 2022). Therefore, we aimed to study the effects of the main compounds of the studied formulation on this enzyme *in silico*. Our findings proved that hesperidin, rutin, and eriodictyol released higher energy than Atorvastatin, suggesting possible potent HMG-CoA reductase inhibitory potential.

The decline in cholesterol concentrations in rats induced by the compounds from the two extracts (*Zingiber officinale* and *Citrus limon*) was systematically analyzed through molecular docking interactions with the primary protein responsible for synthesizing various cholesterol types, specifically “HMG-CoA reductase”. This protein has long been recognized as a pivotal receptor targeted by compounds to regulate body homeostasis (Istvan and Deisenhofer, 1997; Libby and Bornfeldt, 2020). The table presents the results, facilitating the identification of molecules exhibiting potential inhibitory characteristics, as evidenced by their comparative values against the native ligand of the specific protein.

The “HMG-CoA reductase” enzyme is pivotal in cholesterol biosynthesis (Istvan and Deisenhofer, 1997; Libby and Bornfeldt, 2020), and substrate concentrations often modulate it. Its significance extends to cellular homeostasis (Bekkouch et al., 2022, 2019). In our investigation, a protein was selected from the protein repository “1DQ9” to probe its interactions with compounds from our extracts. Notably, three molecules exhibited binding affinities comparable to or surpassing those of the native ligand, Atorvastatin.

Rutin demonstrated a commendable binding affinity with a score of (-9 kcal/mol), establishing robust intramolecular interactions within the enzyme’s active site. These included hydrogen bonding patterns with key residues such as LYS C:633 and HIS C:536. Concurrently, other interaction motifs were evident, encompassing alkyl and pi-alkyl configurations and pi-cationic interactions.

With a score of -10.4 kcal/mol, hesperidin forged significant hydrogen bonds with pivotal residues across chains C and B. The intricate network of interactions also encompassed alkyl, pi-alkyl, pi-cationic, and specific unfavorable donor-donor motifs.

Lastly, eriodictyol, with a score of (-8.5 kcal/mol), manifested notable interactions, including hydrogen bonding and other distinct patterns, delineating its potential binding modes within the protein’s architecture.”The production of cholesterol is regulated by HMG-CoA reductase, which acts as the factor that limits the rate of the mevalonate pathway (Ezeh and Ezeudemba, 2021b). Inhibiting the activity of this enzyme is one of the most effective treatments for reducing LDL levels, thereby preventing cardiovascular disease and atherosclerosis. (Charan et al., 2022). Therefore, we aimed to study the effects of the main compounds of the studied formulation on this enzyme in silico. Our findings proved that hesperidin, rutin, and eriodictyol released higher energy than Atorvastatin, suggesting possible potent HMG-CoA reductase inhibitory potential.

The remaining molecules exhibited notable energy release; however, it was surpassed by the reference molecule, Atorvastatin.

These findings support the remarkable hypocholesterolemic effect revealed in the *in vivo* study. These molecules can be the subject of several in-depth studies to use as therapeutic agents against cholesterol increase.

The findings from both the acute and subacute toxicity studies demonstrate that the tested extracts are safe and non-toxic. No adverse effects were observed in the acute toxicity evaluation, even at the highest administered doses. The animals showed no distress, behavioral changes, or mortality, reflecting the extracts’ high safety margin. Similarly, the subacute toxicity results reinforce this conclusion. The organ weight analysis revealed that absolute and relative liver and kidney weights remained stable across all treatment groups, regardless of the dose or type of extract (ginger juice, lemon juice, or their combined formulation). This stability suggests that the extracts caused no harm to these vital organs. Moreover, the hematological parameters, including white and red blood cell counts, hemoglobin levels, hematocrit, platelet counts, and erythrocyte indices, were consistent with normal physiological ranges and comparable to the control group. Together, these results provide compelling evidence that both acute and subacute administrations of the extracts are well-tolerated and do not induce toxicity, underscoring their potential safety for therapeutic use.

The comparative analysis of our molecular docking results with those reported in the three referenced articles highlights key similarities and differences in the interactions of natural compounds with HMG-CoA reductase, a critical enzyme in cholesterol biosynthesis. Our study demonstrated that hesperidin and rutin, derived from *Zingiber officinale* and *Citrus limon*, exhibit binding energies of -10.4 kcal/mol and -9.0 kcal/mol, respectively, reflecting strong inhibitory potential. These results are consistent with the findings of Medina-Franco et al. (Medina-Franco et al., 2005), who reported significant interactions for α-asarone (-6.60 kcal/mol), mediated by hydrogen bonds with key residues Lys-691 and Glu-559. Similarly, Lin et al. identified curcumin and salvianolic acid C as effective inhibitors, with binding energies complemented by IC_50_ values of 4.3 µM and 8 µM, showcasing their dual computational and experimental validation (Lin et al., 2015).

Jensi et al. provided additional insights by evaluating quercetin and leucocyanidin, demonstrating stable hydrogen-bonding interactions with HMG-CoA reductase, significantly reducing LDL cholesterol levels in treated animal models. While our results for 6-gingerol (-6.1 kcal/mol) reveal moderate binding energy, the superior affinities of hesperidin and rutin underscore their enhanced therapeutic potential compared to quercetin and leucocyanidin. This comparative finding highlights the robust inhibitory activity of flavonoids and suggests their potential for further optimization in drug design (Dhivya Jensi and Ananda Gopu, 2018).

What sets our study apart is the demonstration of synergistic effects when combining ginger and lemon extracts, which was not extensively explored in the referenced articles. This combination yielded enhanced binding affinities and superior hypolipidemic outcomes, including more significant reductions in LDL cholesterol and increases in HDL cholesterol levels compared to the effects of individual compounds. These findings underscore the value of exploring synergistic formulations to maximize therapeutic efficacy. Together, these insights provide compelling evidence for developing natural-product-based therapies as safer and potentially more effective alternatives to synthetic statins, warranting further investigation through pre-clinical and clinical studies.

Based on these data and our findings, the *Zingiber officinale* and *Citrus limon* juice formulations contain flavonoids and gingerols that exert a synergetic lipid-lowering effect by regulating lipid metabolism and are entirely safe.

## 5. Conclusions

This study evaluated the lipid-lowering effect of *Zingiber officinale* and *Citrus limon* juices and their combination in triton-induced hyperlipidemic mice. Moreover, we carried out an *in silico* study to support the anti-hyperlipidemic effect of these treatments. The findings revealed that the juice of both plants exerted significant lipid-lowering effects. Interestingly, the combination of the juices revealed higher lipid-lowering effects, suggesting the synergetic activity of the bioactive phytochemicals of *Zingiber officinale* with those of *Citrus limon*. Moreover, the *in silico* study showed that some of the main compounds of *Citrus limon*, which are eriodictyol, rutin, hesperidin, and isorhamnetin, could have a higher potential to inhibit the activity of the HMG-CoA reductase enzyme. We conclude that *Zingiber officinale* and *Citrus limon* juices and their formulation could be potent natural alternative treatments for hyperlipidemia. As a perspective for the work, we could consider exploring the potential clinical applications of *Zingiber officinale* and *Citrus limon* extracts in the treatment of chronic hyperlipidemia while deepening our understanding of the underlying molecular mechanisms and optimizing their integration into effective phytotherapeutic formulations. By addressing these research gaps, our findings provide a holistic perspective on the multifaceted approach to managing acute hyperlipidemia, offering a foundation for future investigations in the field.

## Acknowledgments

The authors of this work wish to express their appreciation to Mrs. Badraoui Mustapha, Ramdaoui Karim, and Joudar Mohammed for their technical assistance and material and moral support.

- This manuscript/data, or parts thereof, has not been submitted for possible publication in another journal.

## Data Availability Statement

All data generated or analyzed during this study are included in this published article

## Ethics approval and consent to participate

Animal housing and experimental procedures followed the European Union directive (2010/63/E.U.). The study was approved by the Faculty of Sciences Institutional Review Board of Oujda University, Morocco (03/21-LBBEH-15 and March 5, 2021).

## Authors contributions

**Conceptualization:** Oussama Bekkouch, Jinwon Choi, and Bonglee Kim**; Methodology:** Oussama Bekkouch, Ayoub Bekkouch, Sojin Kang, Sojin Kang, Bonglee Kim, Mohammed Bourhia, Hamza Elbouny, and Souliman Amrani**.; Validation:** Mohammed Choukri, Bonglee Kim, and Souliman Amrani**; Formal analysis:** Ilham Touiss, Soufiane El Asri, Min Choi, Chin-Hoon Ahn, and Kiryang Kim**; Resources:** Hyo Jeong Kim**; Data curation:** Ayoub Bekkouch**.; Writing—original draft preparation:** Oussama Bekkouch, Ayoub Bekkouch, and Bonglee Kim**.; Writing—review and editing:** Oussama Bekkouch, Moon Nyeo Park**; Supervision:** Mohammed Choukri, Bonglee Kim, and Souliman Amrani**; Funding acquisition:** Bonglee Kim. All authors have read and agreed to the published version of the manuscript.

## Plant collection approval

No approval is needed for *Zingiber officinale* and *Citrus limon* research purposes.

## IUCN Policy statement

Plant material collection complies with relevant institutional, national, and international guidelines and legislation.

## Abbreviations

The following abbreviations are used in this manuscript:

ABTS: 2,2-azino-bis (3-ethylbenzothiazoline-6-sulfonic acid)]
AIP: Atherogenic index of plasma
ATG: Atorvastatin-treated group
BHA: Butylated hydroxy anisole
CHOL: Cholesterol
DPPH: 1,1-Diphenyl picryl hydrazil
DW: Dry weight
FTG: Formulation-treated group
GAE: Gallic acid equivelent
GJ: Ginger juice
GJTG: Ginger juice-treated group
HCG: Hypercholesterolemic group
HCT: Hematocrit
HDL: High-density lipoprotein
HFD: High-fat diet
HGB: Hemoglobin concentration
HPLC: High-performance liquid chromatography
IC50: Inhibitory concentration 50
LDL: Low-density lipoprotein
LJ: Lemon juice
LJTG: Lemon juice-treated group
MCH: Mean corpuscular hemoglobin
MCHC: Mean corpuscular hemoglobin concentration
MCV: Mean corpuscular volume
MDA: Malondialdehydes
MPV: Mean platelet volume
NCG: Normocholesterolemic group
PLT: Platelets
QE: Quercetin equivalent
RBC: Red blood cells
TAC: Total antioxidant capacity
TBARS: Thiobarbituric acid species
TC: Total cholesterol
TG: Triglycerides
VLDL: Very low-density lipoprotein
WBC: White blood cells

## References

Abdou, H.M., Yousef, M.I., Newairy, A.A., 2018. Triton WR-1339-induced hyperlipidemia, DNA fragmentation, neurotransmitters inhibition, oxidative damage, histopathological and morphometric changes: the protective role of soybean oil. The Journal of Basic and Applied Zoology 79. 10.1186/s41936-018-0065-z

Ainsworth, E.A., Gillespie, K.M., 2007. Estimation of total phenolic content and other oxidation substrates in plant tissues using Folin – Ciocalteu reagent. Nat Protoc 2, 875–877. 10.1038/nprot.2007.102

Alberti, K.G.M.M., Zimmet, P., Shaw, J., 2005. The metabolic syndrome - A new worldwide definition. Lancet 366, 1059–1062. 10.1016/S0140-6736(05)67402-8

Ali, B.H., Blunden, G., Tanira, M.O., Nemmar, A., 2008. Some phytochemical, pharmacological and toxicological properties of ginger (Zingiber officinale Roscoe): A review of recent research. Food and Chemical Toxicology 46, 409–420. 10.1016/j.fct.2007.09.085

Ali, S.M., Chen, P., Sheikh, S., Ahmad, A., Ahmad, M., Paithankar, M., Desai, B., Patel, P., Khan, M., Chaturvedi, A., Patel, R., Panchal, D.T., Shah, K., Chavda, V., Saboo, B.D., Patel, A., Ahmad, I., 2021. Thymoquinone with Metformin Decreases Fasting, Post Prandial Glucose, and HbA1c in Type 2 Diabetic Patients. Drug Res 71, 302–306. 10.1055/a-1388-5415

Ama Moor, V.J., Nya Biapa, P.C., Nono Njinkio, B.L., Moukette Moukette, B., Sando, Z., Kenfack, C., Ateba, B., Ngo Matip, M.E., Pieme, C.A., Ngogang, J., 2017. Hypolipidemic effect and activation of Lecithin Cholesterol Acyl Transferase (LCAT) by aqueous extract of Spirulina platensis during toxicological investigation. BMC Nutr 3, 1–8. 10.1186/s40795-017-0146-2

Amorim, J.L., Simas, D.L.R., Pinheiro, M.M.G., Moreno, D.S.A., Alviano, C.S., Da Silva, A.J.R., Fernandes, P.D., 2016. Anti-inflammatory properties and chemical characterization of the essential oils of four Citrus species. PLoS One 11, 1–18. 10.1371/journal.pone.0153643

Antoniewicz (Kałduńska), J., Jakubczyk, K., Gutowska, I., Janda, K., 2021. Ginger (Zingiber officinale) – a spice with therapeutic properties. Medycyna Ogólna i Nauki o Zdrowiu 27, 40–44. 10.26444/monz/134013

Auer, J., Sinzinger, H., Franklin, B.A.R.R.Y., Berent, R., 2016. Muscle- and skeletal-related side-effects of statins: Tip of the iceberg? Eur J Prev Cardiol 23, 88–110. 10.1177/2047487314550804

Bekkouch, O., Dalli, M., Harnafi, M., Touiss, I., Mokhtari, I., El Assri, S., Harnafi, H., Choukri, M., Ko, S.J., Kim, B., Amrani, S., 2022. Ginger (Zingiber officinale Roscoe), Lemon (Citrus limon L.) Juices as Preventive Agents from Chronic Liver Damage Induced by CCl4: A Biochemical and Histological Study. Antioxidants 11. 10.3390/antiox11020390

Bekkouch, O., Harnafi, M., Touiss, I., Khatib, S., Harnafi, H., Alem, C., Amrani, S., 2019. In Vitro Antioxidant and in Vivo Lipid-Lowering Properties of Zingiber officinale Crude Aqueous Extract and Methanolic Fraction: A Follow-Up Study. Evidence-based Complementary and Alternative Medicine 2019. 10.1155/2019/9734390

Bouhlali, E. dine T., Hmidani, A., Bourkhis, B., Khouya, T., Harnafi, H., Filali-Zegzouti, Y., Alem, C., 2020. Effect of Phoenix dactylifera seeds (dates) extract in triton WR-1339 and high-fat diet induced hyperlipidaemia in rats: A comparison with simvastatin. J Ethnopharmacol 259. 10.1016/j.jep.2020.112961

Brahma Naidu, P., Uddandrao, V.V.S., Ravindar Naik, R., Suresh, P., Meriga, B., Begum, M.S., Pandiyan, R., Saravanan, G., 2016. Ameliorative potential of gingerol: Promising modulation of inflammatory factors and lipid marker enzymes expressions in HFD induced obesity in rats. Mol Cell Endocrinol 419, 139–147. 10.1016/j.mce.2015.10.007

Cha, J.H., Kim, S.R., Kang, H.J., Kim, M.H., Ha, A.W., Kim, W.K., 2016. Corn silk extract improves cholesterol metabolism in C57BL/6J mouse fed high-fat diets. Nutr Res Pract 10, 501–506. 10.4162/nrp.2016.10.5.501

Chaachouay, N., Azeroual, A., Bencharki, B., Zidane, L., 2022. Herbal medicine used in the treatment of cardiovascular diseases in the Rif, North of Morocco. Front Pharmacol 13. 10.3389/fphar.2022.921918

Charan, J., Riyad, P., Ram, H., Purohit, A., Ambwani, S., Kashyap, P., Singh, G., Hashem, A., Abd-Allah, E.F., Gupta, V.K., Kumar, A., Panwar, A., 2022. Ameliorations in dyslipidemia and atherosclerotic plaque by the inhibition of HMG-CoA reductase and antioxidant potential of phytoconstituents of an aqueous seed extract of Acacia senegal (L.) Willd in rabbits. PLoS One 17. 10.1371/journal.pone.0264646

Chen, G.-L., Chen, S.G., Xie, Y.Q., Chen, F., Zhao, Y.Y., Luo, C.X., Gao, Y.Q., 2015. Total phenolic, flavonoid and antioxidant activity of 23 edible flowers subjected to in vitro digestion. J Funct Foods 17, 243–259. 10.1016/j.jff.2015.05.028

Chen, W., S.H., X.Y., & H.Y., 2018. Antioxidant interactions between polyphenol-rich plant extracts: A review of mechanisms and applications. Mol Nutr Food Res 55, 312–329.

Cockcroft, S., 2021. Mammalian lipids: Structure, synthesis and function. Essays Biochem. 10.1042/EBC20200067

Dhivya Jensi, V., Ananda Gopu, P., 2018. Comparative Analysis of Quercetin and Leucocynidin against HMG-CoA reductase and their evaluation of hypolipidemic activity. 10.2174/1573401314666181228121042

Dujovne, C.A., n.d. Side Effects of Statins: Hepatitis Versus “Transaminitis”-Myositis Versus “CPKitis.”

El Moussaoui, A., Jawhari, F.Z., Almehdi, A.M., Elmsellem, H., Fikri Benbrahim, K., Bousta, D., Bari, A., 2019. Antibacterial, antifungal and antioxidant activity of total polyphenols of Withania frutescens.L. Bioorg Chem. 10.1016/j.bioorg.2019.103337

Elbouny Hamza, O.B.S.K.A.Chakib., 2023. Hypolipidemic Effect of Thymus munbyanus subsp. ciliatus Greuter & Burdet.: Guinea Pig as a Model for Tyloxapol-induced Hyperlipidemia. Journal of Biologically Active Products from Nature 12, 507–513.

Enstitüsü, B.A.T.A., 2015. Growing And Plant Characteristics Of Ginger (Zingiber officinale Roscoe). Food and Agiculture Organization 22, 1–9.

Ezeh, K.J., Ezeudemba, O., 2021a. Hyperlipidemia: A Review of the Novel Methods for the Management of Lipids. Cureus. 10.7759/cureus.16412

Ezeh, K.J., Ezeudemba, O., 2021b. Hyperlipidemia: A Review of the Novel Methods for the Management of Lipids. Cureus. 10.7759/cureus.16412

Farkhondeh, T., Samarghandian, S., Pourbagher-Shahri, A.M., 2019. Hypolipidemic effects of Rosmarinus officinalis L. J Cell Physiol 234, 14680–14688. 10.1002/jcp.28221

Farnier, M., Zeller, M., Masson, D., Cottin, Y., 2021. Triglycerides and risk of atherosclerotic cardiovascular disease: An update. Arch Cardiovasc Dis 114, 132–139. 10.1016/j.acvd.2020.11.006

Friedman, M., Byers, S.O., 1953. The mechanism responsible for the hypercholesteremia induced by triton WR-1339. J Exp Med 97, 117–130. 10.1084/jem.97.1.117

Ghatak, S.B., Panchal, S.J., 2012. Anti-hyperlipidemic activity of oryzanol, isolated from crude rice bran oil, on triton WR-1339-induced acute hyperlipidemia in rats. Brazilian Journal of Pharmacognosy 22, 642–648. 10.1590/S0102-695X2012005000023

Gong, G., Guan, Y.Y., Zhang, Z.L., Rahman, K., Wang, S.J., Zhou, S., Luan, X., Zhang, H., 2020. Isorhamnetin: A review of pharmacological effects. Biomedicine and Pharmacotherapy 128. 10.1016/j.biopha.2020.110301

Hamideh Parhiz, Ali Roohbakhsh, Fatemeh Soltani, Ramin Rezaee, M.I., 2014. Antioxidant and Anti-Inflammatory Properties of the Citrus Flavonoids Hesperidin and Hesperetin: An Updated Review of their Molecular Mechanisms and Experimental Models. Phytotherapy Research 29.

Himed, L., Merniz, S., 2016. Algerian Journal of Natural Products essentielle de Citrus limon ( variété Lisbon) extraite par hydrodistillation 1, 252–260.

Ibrahim, A.Y., Hendawy, S.F., Elsayed, A.A.A., Omer, E.A., 2016. Evaluation of hypolipidemic Marrubium vulgare effect in Triton WR-1339-induced hyperlipidemia in mice. Asian Pac J Trop Med 9, 453–459. 10.1016/j.apjtm.2016.03.038

Istvan, E.S., Deisenhofer, J., 1997. The structure of the catalytic portion of human HMG-CoA reductase.

Jaiswal, S.K., Gupta, V.K., Siddiqi, N.J., Pandey, R.S., Sharma, B., 2015. Hepatoprotective Effect of Citrus limon Fruit Extract against Carbofuran Induced Toxicity in Wistar Rats. Chinese Journal of Biology 2015, 1–10. 10.1155/2015/686071

Kaur, C., Kapoor, H.C., 2001. Antioxidants in fruits and vegetables - the millennium’s health. Int J Food Sci Technol 36, 703–725.

Kehal, 2013. Utilisation de l’huile essentielle de Citrus limon comme agent conservateur et aromatique dans la crème fraîche, Mémoire de diplôme de Magister en Sciences alimentaires, Institut de la Nutrition, de l’Alimentation et des Technologies Agro-Alimentaires, Université Constantine 1, Algerie.

Khanna, A.K., Rizvi, F., Chander, R., 2002. Lipid lowering activity of Phyllanthus niruri in hyperlipemic rats. J Ethnopharmacol 82, 19–22. 10.1016/S0378-8741(02)00136-8

Khouya, T., Ramchoun, M., Hmidani, A., Bouhlali, E. dine T., Amrani, S., Alem, C., 2021. Phytochemical analysis and bioactivity evaluation of Moroccan Thymus atlanticus (Ball) fractions. Sci Afr 11. 10.1016/j.sciaf.2021.e00716

Kim, K.J., Ko, S.K., 2017. The accelerating action of lipid excretion of immature Citrus fruits. Korean Journal of Pharmacognosy 48, 134–140.

Kumar, S., P.A.K., & G.R.K., 2019. Antioxidant activity and phenolic content of Citrus limon: A comprehensive review. Curr Nutr Food Sci 22, 1–20. 10.2174/1573401314666181228121042

Li, Y., Tran, V.H., Duke, C.C., Roufogalis, B.D., 2012. Preventive and protective properties of zingiber officinale (Ginger) in diabetes mellitus, diabetic complications, and associated lipid and other metabolic disorders: A brief review. Evidence-based Complementary and Alternative Medicine 2012. 10.1155/2012/516870

Libby, P., Bornfeldt, K.E., 2020. How Far We Have Come, How Far We Have Yet to Go in Atherosclerosis Research. Circ Res 126, 1107–1111. 10.1161/CIRCRESAHA.120.316994

Lin, S.H., Huang, K.J., Weng, C.F., Shiuan, D., 2015. Exploration of natural product ingredients as inhibitors of human HMG-CoA reductase through structure-based virtual screening. Drug Des Devel Ther 9, 3313– 3324. 10.2147/DDDT.S84641

Liu, Z.L., Liu, J.P., Zhang, A.L., Wu, Q., Ruan, Y., Lewith, G., Visconte, D., 2011. Chinese herbal medicines for hypercholesterolemia. Cochrane Database Syst Rev CD008305. 10.1002/14651858.CD008305.pub2

Luo, Y., Sun, G., Dong, X., Wang, M., Qin, M., Yu, Y., Sun, X., 2015. Isorhamnetin attenuates atherosclerosis by inhibiting macrophage apoptosis via PI3K/ AKT activation and HO-1 induction. PLoS One 10, 1–19. 10.1371/journal.pone.0120259

M, A.M., Mastoi, S.M., Saleem, A., Niaz, K., Hakro, S., 2019. Shogaols exhibit cardiodepressant activity at low doses and cardiotonic properties at higher doses 88–90. 10.15406/jccr.2019.12.00441

Manzocco, L., Anese, M., Nicoli, M.C., 1998. Antioxidant Properties of Tea Extracts as Affected by Processing.

Manzoni, A.G., Passos, D.F., da Silva, J.L.G., Bernardes, V.M., Bremm, J.M., Jantsch, M.H., de Oliveira, J.S., Mann, T.R., de Andrade, C.M., Leal, D.B.R., 2019. Rutin and curcumin reduce inflammation, triglyceride levels and ADA activity in serum and immune cells in a model of hyperlipidemia. Blood Cells Mol Dis 76, 13–21. 10.1016/j.bcmd.2018.12.005

Mbikay, M., Sirois, F., Simoes, S., Mayne, J., Chrétien, M., 2014. Quercetin-3-glucoside increases low-density lipoprotein receptor (LDLR) expression, attenuates proprotein convertase subtilisin/kexin 9 (PCSK9) secretion, and stimulates LDL uptake by Huh7 human hepatocytes in culture. FEBS Open Bio 4, 755– 762. 10.1016/j.fob.2014.08.003

Medina-Franco, J.L., López-Vallejo, F., Rodríguez-Morales, S., Castillo, R., Chamorro, G., Tamariz, J., 2005. Molecular docking of the highly hypolipidemic agent α-asarone with the catalytic portion of HMG-CoA reductase. Bioorg Med Chem Lett 15, 989–994. 10.1016/j.bmcl.2004.12.046

Mishra, P. R., Panda, P. K., Apanna, K. C., & Panigrahi, S., 2011. Evaluation of acute hypolipidemic activity of different plant extracts in Triton WR-1339 induced hyperlipidemia in albino rats. Pharmacologyonline 3, 925–934.

Munasinghe, J., Seneviratne, C.K., Thabrew, M.I., Ajith, M., 2001. Antiradical and Antilipoperoxidative Effects of Some Plant Extracts used by Sri Lankan Cardioprotection. Phytotherapy Research 15, 519–523.

Murad, S., Niaz, K., Aslam, H., 2018. Effects of Ginger on LDL-C, Total Cholesterol and Body Weight. Clinical & Medical Biochemistry 04, 4–6. 10.4172/2471-2663.1000140

NR, L., Ravindran Nair, A., Senthilpandian, S., Ravi, V., 2021. Hypolipidemic action of Rutin on Triton WR-1339 induced hyperlipidemia in rats. Journal of Pre-Clinical and Clinical Research 15, 51–55. 10.26444/jpccr/136231

Oboh, G., Bello, F.O., Ademosun, A.O., Akinyemi, A.J., Adewuni, T.M., 2015. Antioxidant, hypolipidemic, and anti-angiotensin-1-converting enzyme properties of lemon (Citrus limon) and lime (Citrus aurantifolia) juices. Comp Clin Path 24, 1395–1406. 10.1007/s00580-015-2088-x

OECD Guideline, 2001. Guidance Document on Acute Oral Toxicity Testing. Guidance document on acute oral toxicity testing.

Pacuła, A.J., Kaczor, K.B., Antosiewicz, J., Długosz, A., Janecka, A., Janecki, T., Wojtczak, A., Ścianowski, J., Santi, C., Bagnoli, L., Dembinski, R., 2017. New chiral ebselen analogues with antioxidant and cytotoxic potential. Molecules 22. 10.3390/molecules22030492

Panahi, Y., Ahmadi, Y., Teymouri, M., Johnston, T.P., Sahebkar, A., 2018. Curcumin as a potential candidate for treating hyperlipidemia: A review of cellular and metabolic mechanisms. J Cell Physiol. 10.1002/jcp.25756

Paul, P., Islam, M.K., Mustari, A., Khan, M.Z.I., 2013. Hypolipidemic Effect of Ginger Extract in Vanaspati Fed Rats. Bangladesh Journal of Veterinary Medicine 10, 93–96. 10.3329/bjvm.v10i1-2.15652

Ramchoun, M., Harnafi, H., Alem, C., Benlyas, M., Elrhaffari, L., Amrani, S., 2009. Study on antioxidant and hypolipidemic effects of polyphenol-rich extracts from Thymus vulgaris and Lavendula multifida. Pharmacognosy Res 1, 106–112.

Rašković, A., Ćućuz, V., Torović, L., Tomas, A., Gojković-Bukarica, L., Ćebović, T., Milijašević, B., Stilinović, N., Cvejić Hogervorst, J., 2019. Resveratrol supplementation improves metabolic control in rats with induced hyperlipidemia and type 2 diabetes. Saudi Pharmaceutical Journal 27, 1036–1043. 10.1016/j.jsps.2019.08.006

Scharwey, M., Tatsuta, T., Langer, T., 2013. Mitochondrial lipid transport at a glance. J Cell Sci 126, 5317– 5323. 10.1242/jcs.134130

Schurr, P.E., Schultz, J.R., Parkinson, T.M., 1972. Triton-induced hyperlipidemia in rats as an animal model for screening hypolipidemic drugs. Lipids 7, 68–74. 10.1007/BF02531272

Shao, Y., Yu, Y., Li, C., Yu, J., Zong, R., Pei, C., 2016. Synergistic effect of quercetin and 6-gingerol treatment in streptozotocin induced type 2 diabetic rats and poloxamer P-407 induced hyperlipidemia. RSC Adv 6, 12235–12242. 10.1039/c5ra16493a

Sharma, I., Cusain, D., Dixit, V.P., 1996. Hypolipidaemic and antiatherosclerotic effects of Zingiber officinale in cholesterol fed rabbits. Phytotherapy Research 10, 517–518. 10.1002/(SICI)1099-1573(199609)10:6<517::AID-PTR839>3.0.CO;2-L

Shen, Y., Song, X., Li, L., Sun, J., Jaiswal, Y., Huang, J., Liu, C., Yang, W., Williams, L., Zhang, H., Guan, Y., 2019. Protective effects of p-coumaric acid against oxidant and hyperlipidemia-an in vitro and in vivo evaluation. Biomedicine and Pharmacotherapy 111, 579–587. 10.1016/j.biopha.2018.12.074

Shende, M., 2024. Investigating the bioactive characteristics of Zingiber officinale and Citrus medica with emphasis on their antioxidant, antimicrobial, and anthelmintic properties. International Journal of Pharmaceutical Research and Development 6, 118–125. 10.33545/26646862.2024.v6.i2b.66

Tiencheu, B., Nji, D.N., Achidi, A.U., Egbe, A.C., Tenyang, N., Tiepma Ngongang, E.F., Djikeng, F.T., Fossi, B.T., 2021. Nutritional, sensory, physico-chemical, phytochemical, microbiological and shelf-life studies of natural fruit juice formulated from orange (Citrus sinensis), lemon (Citrus limon), Honey and Ginger (Zingiber officinale). Heliyon 7, e07177. 10.1016/j.heliyon.2021.e07177

Tirawanchai, N., Supapornhemin, S., Somkasetrin, A., Suktitipat, B., Ampawong, S., 2018. Regulatory effect of Phikud Navakot extract on HMG-CoA reductase and LDL-R: Potential and alternate agents for lowering blood cholesterol. BMC Complement Altern Med 18, 1–8. 10.1186/s12906-018-2327-1

Tohma, H., Gülçin, İ., Bursal, E., Gören, A.C., Alwasel, S.H., Köksal, E., 2017. Antioxidant activity and phenolic compounds of ginger (Zingiber officinale Rosc.) determined by HPLC-MS/MS. Journal of Food Measurement and Characterization 11, 556–566. 10.1007/s11694-016-9423-z

Touiss, I., Khatib, S., Bekkouch, O., Amrani, S., Harnafi, H., 2017. Phenolic extract from Ocimum basilicum restores lipid metabolism in Triton WR-1339-induced hyperlipidemic mice and prevents lipoprotein-rich plasma oxidation. Food Science and Human Wellness 6, 28–33. 10.1016/j.fshw.2017.02.002

www.pymol.org, n.d. PyMOL by Schrödinger [WWW Document].

www.vina.scripps.edu, n.d. AutoDock Vina.

Zhang, H., L.H., & W.Q., 2020. Synergistic effects of ginger and citrus extracts on antioxidant and antimicrobial activities. ournal of Functional Foods 65.

